# Imaging of cellular dynamics *in vitro* and *in situ*: from a whole organism to sub-cellular imaging with self-driving, multi-scale microscopy

**DOI:** 10.1101/2024.02.28.582579

**Authors:** Stephan Daetwyler, Hanieh Mazloom-Farsibaf, Felix Y. Zhou, Dagan Segal, Etai Sapoznik, Jill M. Westcott, Rolf A. Brekken, Gaudenz Danuser, Reto Fiolka

## Abstract

Many biological processes span multiple time and length scales, including developmental processes and cancer metastasis. While light-sheet fluorescence microscopy (LSFM) has become a fast and efficient method for imaging of organisms, cells and sub-cellular dynamics, simultaneous observations across these scales have remained challenging. Moreover, continuous high-resolution imaging inside living organisms has mostly been limited to few hours as regions of interest quickly move out of view due to sample movement and growth. Here, we present a self-driving, multi-resolution light-sheet microscope platform controlled by a custom Python-based software, to simultaneous observe and quantify sub-cellular dynamics and entire organisms *in vitro* and *in vivo* over hours of imaging. We apply the platform to the study of developmental processes, cancer invasion and metastasis, and we provide quantitative multi-scale analysis of immune-cancer cell interactions in zebrafish xenografts.

## Introduction

Over the past two decades, light-sheet microscopy has emerged as a powerful approach for fast and efficient fluorescence imaging^1–3^. At the sub-cellular level, high-resolution light-sheet microscopes such as lattice light-sheet microscopes^4^, field synthesis^5^ , dual-view inverted selective plane illumination microscopes^6^ or axially swept light-sheet microscopes (ASLM)^7,8^ enable studies of spatiotemporal organization and dynamics of cell signaling, cell morphology, and local cell-cell interactions. At the organismal scale, low-resolution light-sheet imaging^9^ provides capabilities to rapidly image several entire organisms over days^10,11^ and study processes such as cell dissemination, migration patterns or to find rare cellular events such as immune-cancer cell interactions. Progress has been made towards combining low- and high-resolution imaging for cleared tissue imaging^12^. However, to our knowledge, no light-sheet imaging platform has been described to dynamically image living organisms in their entirety over several hours while simultaneously imaging at sub-cellular resolution to study biological processes across scales.

The lack of technologies for imaging a whole, living organism at sub-cellular resolution is due to technical and practical limitations^13^. There is a trade-off in optical design between numerical aperture (NA) and working distance as well as field of view^14^, i.e., the higher the imaging resolution, the smaller the region to image (Fig. 1a). From a data collection perspective, there are practical limitations as imaging an entire organism such as a zebrafish larva with sub-cellular resolution would require several hours, precluding any dynamic time-lapse imaging. Moreover, proper Nyquist sampling of the 3D sample space would result in enormous data sizes, e.g., over half a Terabyte per timepoint and channel (Fig. 1b). To overcome these trade-offs, novel, adaptive imaging schemes are required that perform high-resolution imaging only in selected regions of interest and timepoints.

**Fig. 1:**
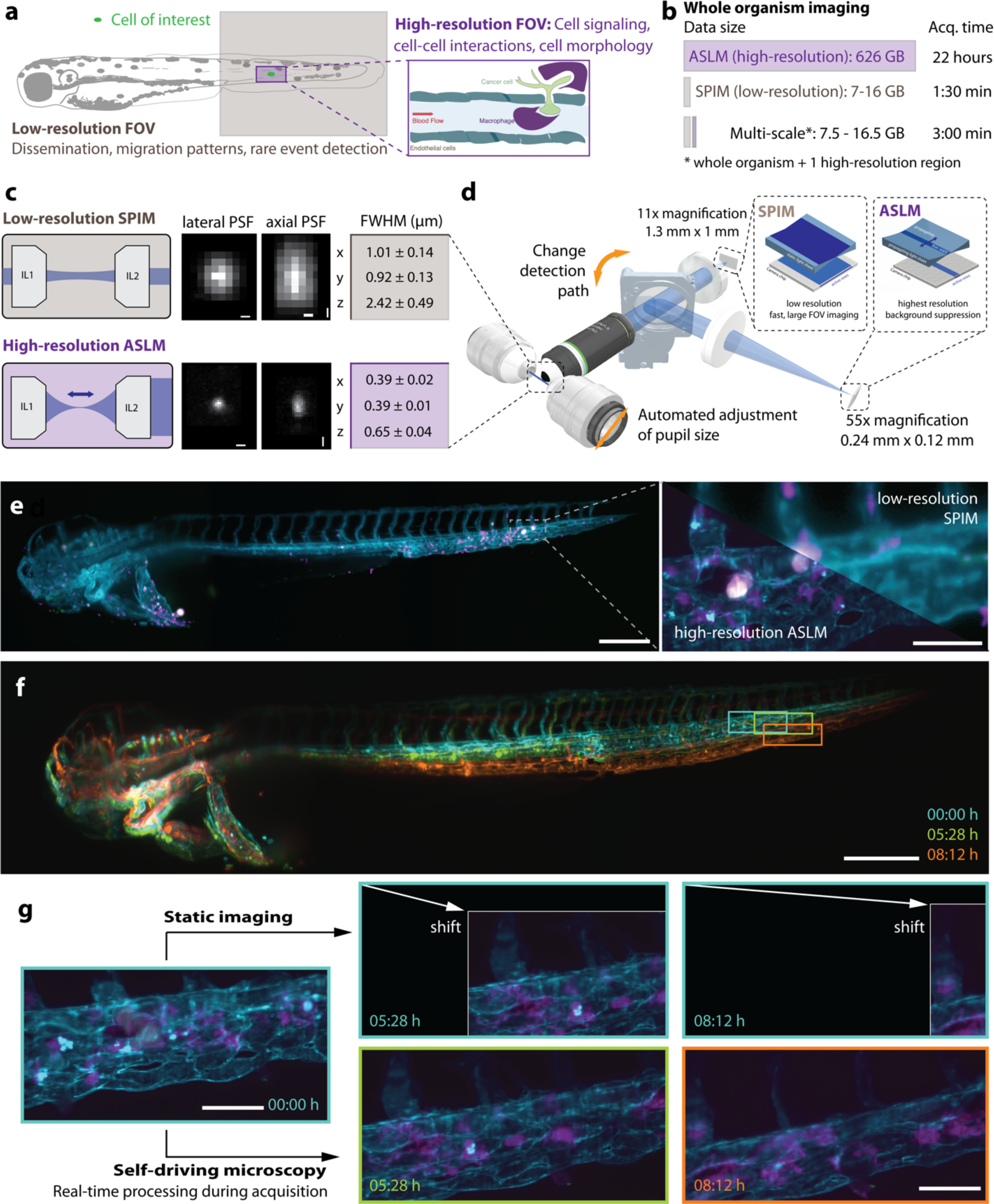
Self-driving, multi-scale microscopy. **a** Schematic of the field of views of low-(brown) and high-(violet) resolution light-sheet imaging modalities to image biological processes at different scales: from sub-cellular to systemic, organismal scales. **b** Calculations of the duration and resulting data sizes for imaging a 2.5-day old zebrafish larva in one fluorescent channel with high-resolution imaging using axially swept light-sheet microscopy (ASLM), low-resolution imaging and a multi-scale imaging approach that captures the whole organisms and one selected region of interest with high-resolution. **c-d** Schematic of our light-sheet microscope platform that enables imaging at low resolution (brown) with dual sided multi-directional selective plane illumination (mSPIM) and at high resolution (violet) with single-sided axially swept light microscopy (ASLM). **c** (Left panels) Schematic light-sheet profiles (top view) of two-sided mSPIM and single-sided ASLM, with the dark blue arrow indicating the scan direction of the thin ASLM light-sheet. (Middle panels) Representative bead profiles of 0.2 *μ*m YG Nanospheres for both modalities. (Right panels) PSF values measured with MetroloJ (n=20 for each modality). **d** Simplified rendering of the multi-scale microscope. The low- and high-resolution modalities share the same illumination and detection objectives. A motorized slit adjusts the effectively used pupil size for illumination to change the light-sheet properties. The detection path is selected with a motorized flip mirror to either the low-resolution mSPIM imaging with a 11.1x magnification and an effective field of view of 1.3 x1 mm^2^, or to high-resolution ASLM imaging with a 55.5x magnification and a typical field of view of 0.24 × 0.12 mm^2^. **e** Representative images from the multi-scale microscope displaying a 2.5 days old zebrafish larva (cyan: vascular marker *Tg(kdrl:Hsa.HRAS-mCherry),* magenta: macrophage marker *Tg(mpeg1:EGFP))* xenografted with U-2 OS osteosarcoma cells (white). Left: Low-resolution image of the whole zebrafish larva. Right: Zoom in to boxed image to compare high-resolution image acquired with ASLM (bottom left) to the mSPIM image (top right). **f** Representative low-resolution stills from a time-lapse imaging experiment of zebrafish larva expressing the vascular marker *Tg(kdrl:Hsa.HRAS-mCherry)* are overlaid onto each other, with the color indicating imaging time (starting at 3 days post fertilization). This highlights sample movement and growth over the observation window and the shift of a region of interest over time (boxed region). **g** High-resolution imaging with a static, pre-defined imaging volume within the microscope coordinate system (top) and self-driving microscopy (bottom). Same color scheme as in **e**. Scale-bar lengths are as follows: **c** 0.5 μm; **e** 250 μm (left), 60 μm (right); **f** 250 μm; **g** 50 μm.

Leveraging adaptive imaging schemes with different imaging modalities, Almada *et al.*^15^ performed unsupervised, event-driven sample treatment and live-to-fixed imaging. In another approach, Alvelid *et al*.^16^ performed automated multiscale imaging using STED super-resolution imaging guided by widefield microscopy. Similarly, smart lattice microscopy has been described for imaging rare cellular processes in a culture dish using epifluorescent imaging for detection^17^. While these approaches integrate information from different modalities, they do not allow for multi-resolution imaging of entire, complex 3D samples and organisms, away from a culture dish, with simultaneous observations of sub-cellular processes of interest *in situ*.

To overcome these limitations, we describe here a novel self-driving, multi-resolution light-sheet microscope platform that enables observation and quantification of biological processes across scales *in vivo* over many hours. To this end, we have modularly combined the strengths of low-resolution static light-sheet imaging^10^ with sub-cellular ASLM imaging^7,8^. Guided by custom hardware control software, the microscope is self-driving, i.e., it adjusts the region for high-resolution imaging on-the-fly as the sample develops, grows, or changes. We demonstrate the power of this self-driving, multi-scale imaging platform for diverse applications *in vitro* and *in vivo*, including imaging of cancer spheroids, developmental processes, and cancer cell metastasis in a zebrafish xenograft model. In depth, we detect, visualize, and quantify immune–cancer cell interactions that lead to successful phagocytosis or immune evasion of human cancer cells in a micro-metastasis niche.

## Results

### Multi-sample, multi-resolution system design

We modularly combined dual sided multi-directional selective plane illumination (mSPIM)^18^ for low resolution imaging and single sided axially swept light-sheet microscopy (ASLM)^7,8^ for high resolution imaging (Fig. 1c, Extended Fig. 1). While mSPIM excels at whole organism imaging to reveal cellular and tissue level processes such as cell spreading and migration patterns, and detect rare cellular processes in large volumes, ASLM provides the necessary resolution enhancement to reveal sub-cellular signaling, cell-cell interactions and local changes of cell morphologies (Fig. 1a, c). Both modalities were combined into the same system using a shared illumination path and the same detection objective (Fig. 1d). A description of all hardware components is available as Supplementary Table 1.

Specifically, for illumination, we applied dual-sided mSPIM to illuminate instantaneously a large field of view for whole organism imaging (Supplementary Fig. 1). For sub-cellular imaging, the ASLM modality employed a thin light-sheet (Supplementary Fig. 2), which was scanned in its propagation direction in a coordinated manner with the rolling shutter on a camera. The scanning of the light-sheet allowed us to extend the otherwise beam-waist limited field of view of a thin light-sheet. To change the light-sheet properties for the two modes, we employed a motorized slit conjugate to the pupil of one of the illumination objectives. Upon opening the slit, the numerical aperture of the light-sheet was increased up to 0.4.

For detection, we determined that using a medium magnification objective (20X Olympus XLUMPLFLN) with a high numerical aperture (NA) of 1.0, a large working distance (2 mm) and a large field of view allowed us to use the same detection path for both modalities. A motorized flip mirror after the detection objective enabled us to image at different magnifications by selecting a different tube lens and camera pair (Fig. 1d). In the low-resolution path, a 100 mm tube lens demagnified the image to 11.1x magnification, while a 500 mm tube lens enabled 55.56x magnification in the high-resolution detection path. Moreover, in the low-magnification, low-resolution detection, we used a camera (Iris 15, Photometrics) with a large sensor size of 5056 × 2960 pixels. This provided a 1.9×1.1 mm^2^ large field of view with a deliberate slight under-sampling (382nm demagnified pixel size) for fast imaging of entire organisms. Due to the field number of the objective, this field of view reduced to an effective field of view of 1.3×1.1 mm^2^. In the high-resolution detection path, we used a sensitive camera (Prime BSI Express, Photometrics) with 2048 × 2048 pixels, enabling Nyquist sampled (117nm demagnified pixel size) high resolution imaging using ASLM with the Photometrics programmable scan mode (Supplementary Fig. 3).

To characterize the optical performance of our multi-scale microscope, we determined the point spread function values of 0.2 *μ*m YG Nanospheres embedded in agarose (Figure 1c, Supplementary Note 2). The ASLM resolution was measured laterally as: 0.39 ± 0.02 μm (x) and 0.39 ± 0.01 μm (y); and axially as: 0.65±0.04 μm. For the low-resolution mode, due to a combination of the thicker light-sheet and lower lateral sampling, we measured a lateral resolution of 1.01 ± 0.14 μm (x), 0.92 ± 0.13 μm (y), and an axial resolution of 2.42 ± 0.49 μm (Fig 1c). These values considerably improve over a recently published multi-scale cleared tissue microscope with 2.91 ± 0.31μm axial PSF in the high-resolution mode and a 5.48 ± 1.08 μm axial PSF in the low-resolution mode^12^. In principle, high-resolution imaging without ASLM is possible, however, this led to a reduced axial resolution (Supplementary Fig. 4).

Our low-resolution imaging mode spanned a 1.3×1.0 mm^2^ field of view, which allowed efficient tiling of an entire zebrafish larva. In contrast, the ASLM mode was applied for targeted sub-cellular imaging and spanned typically a field of view of 0.24 × 0.12 mm^2^. Applying both microscope modes enabled efficient imaging of large volumes in the low-resolution mode complemented with a high-resolution modality to zoom in to selected regions of interest for sub-cellular imaging (Fig. 1e). Importantly, the time to switch between both modalities was only limited by the time it took for the flip mirror and the motorized slit to change the light-sheet properties.

### Multi-scale control software

To control all aspects of multi-scale imaging, we implemented our own hardware control software. The software is designed for easy integration of all hardware components and image processing algorithms, inclusion of features for simultaneous acquisition and data processing, handling of all multi-scale aspects such as imaging modality selection, and with a clear model-view-controller (MVC) design pattern that enables future transition of the software to different microscope hardware.

We implemented the software in Python, a popular choice for microscope control software solutions^19,20^. Precise timing of all hardware components was ensured with Python-based control of an NI Data Acquisition card (NI PCIe-6738) with sufficient digital and analog outputs (Supplementary Fig. 6). Importantly, Python offers many optimized image processing libraries such as OpenCV^21^ that can be easily integrated into a Python based hardware control software for automated microscope control. To enable simultaneous processing and acquisition of image data, we leveraged concurrency tools from Thayer, York *et al*.^22^ such as shared memory arrays, threads, and subprocesses. Briefly, these tools span child processes to make hardware control IO-limited rather than CPU-limited. Moreover, by using a buffer queue of shared memory arrays, acquired images could be processed, while newly acquired images were saved to a different shared memory array. Furthermore, the MVC design separated the GUI from the controller and the model, including the routines for on-the-fly image processing, hardware control and concurrency to improve the logic (Supplementary Fig. 7). Using Napari^23^, we further implemented a state-of-the art and open source image viewer to visualize acquired images (Supplementary Fig. 8). The whole control software is available on GitHub: https://github.com/DaetwylerStephan/selfdriving-multiscale-imaging.

### Self-driving multi-scale microscopy

While longitudinal imaging by LSFM over days has been previously performed at cellular resolution^10,11^, high resolution imaging (i.e. at sub-cellular resolution) inside a living organism has so far been restricted to a few hours^24,25^. This limitation arises as a (small) selected region of interest for high-resolution imaging will quickly move out of view due to non-linear sample growth and changes, sample movement, and hardware drift (Fig 1f). Additionally, cells of interest might migrate within the organism away from the selected region. As mentioned previously, this cannot be simply remedied by imaging a larger volume at higher resolution, due to the correspondingly higher sampling requirements (Fig. 1b).

To overcome these challenges and image a selected region of interest over hours, we implemented “self-driving microscopy” within our software (Fig. 1g). We leveraged the low-resolution information of the microscope to guide the acquisition of the high-resolution region using a custom 3D region tracking algorithm (Extended Fig. 2). To successfully apply region tracking on-the-fly, rapid image processing is paramount. Moreover, the tracking should be independent of the content of the high-resolution region so that any fluorescent channel or even brightfield transmission imaging (Extended Fig. 3) could be used to automatically guide high-resolution acquisitions. Thereby, most of the fluorescent photon budget remains available for high-resolution imaging.

**Fig. 2.**
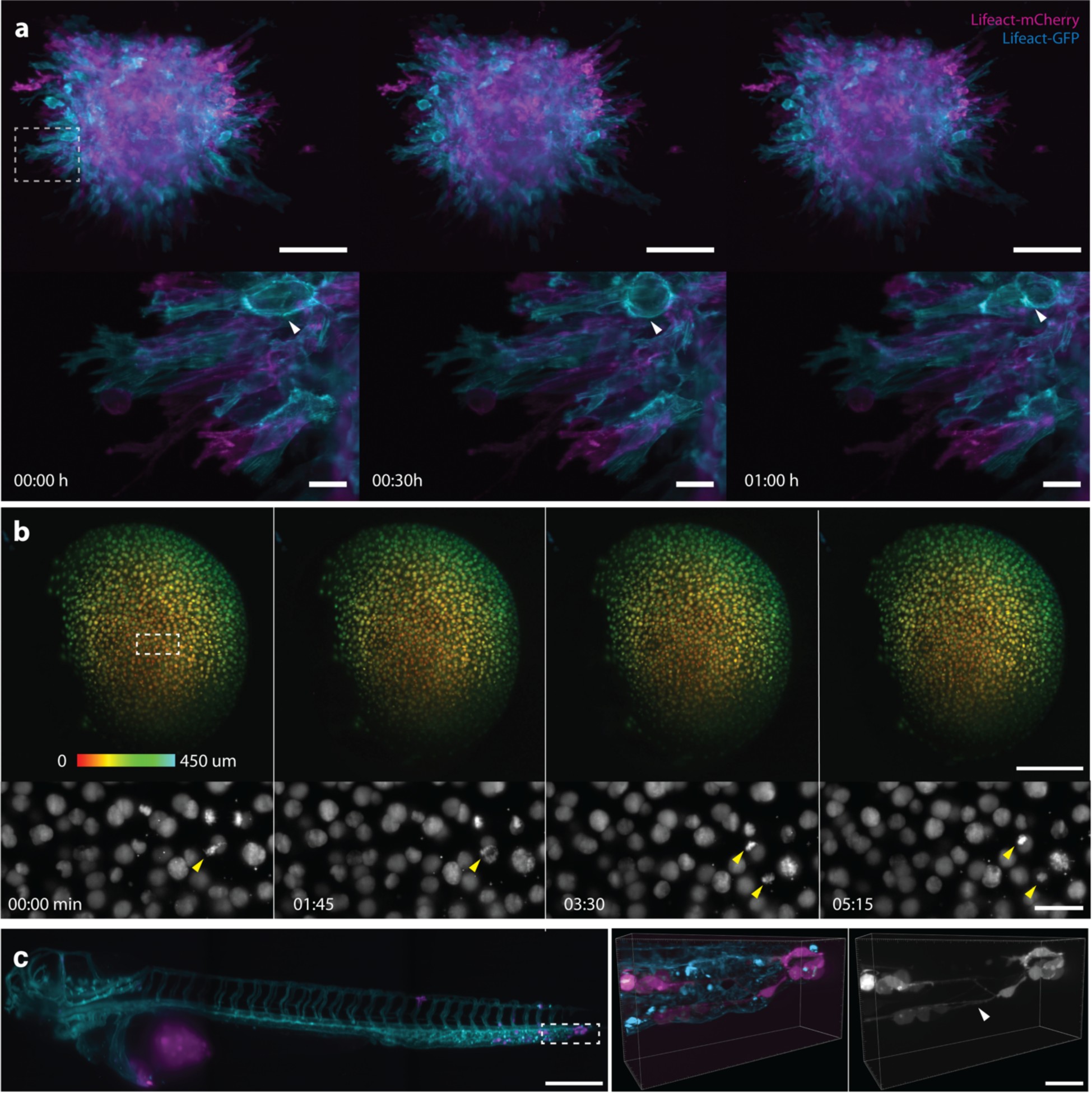
Applications of self-driving, multi-scale microscopy. **a** Multi-scale imaging of a SUM 159 breast cancer cell spheroid embedded into a collagen matrix (Supplementary Movie 1) with the low-resolution modality shown on top and the high-resolution modality on the bottom. The spheroids consisted of a 1:1 mixture of cells expressing the actin marker Lifeact-GFP (cyan) and Lifeact-mCherry (magenta). Boxed region on top indicates the location of the high-resolution region on the bottom. The white arrowhead points at a cell division at the invasive front. **b** Multi-scale imaging of zebrafish gastrulation with cells expressing the histone marker *Tg(h2afva:h2afva-GFP)*, starting at around 6 hours post fertilization (Supplementary Movie 2). Low-resolution imaging (on top) captured the entire zebrafish embryo (color scale: depth of data in 3D volume from 0 to 450 um), while the high-resolution imaging (bottom) enabled near-simultaneous imaging of cell division (yellow arrowheads), including sister chromatid separations. **c** Multi-scale imaging of human breast cancer cells MDA-MB-231 expressing F-tractin-EGFP (left: magenta, right: grayscale) *in situ* in a larval zebrafish xenograft model (Supplementary Movie 3). The cancer cells were xenografted at 2.25 days post fertilization into a zebrafish larvae expressing the vascular marker *Tg(kdrl:Hsa.HRAS-mCherry)* (cyan). Cell spreading patterns as imaged in the low-resolution mode are shown on the left, and a 3D rendering of the high-resolution data is shown on the right. The white arrowhead points at a network of protrusions. Scale-bar lengths are as follows: **a** 150 μm (top), 20 μm (bottom); **b** 200 μm (top), 30 μm (bottom); **b** 300 μm (left); 40 μm (right).

**Fig. 3:**
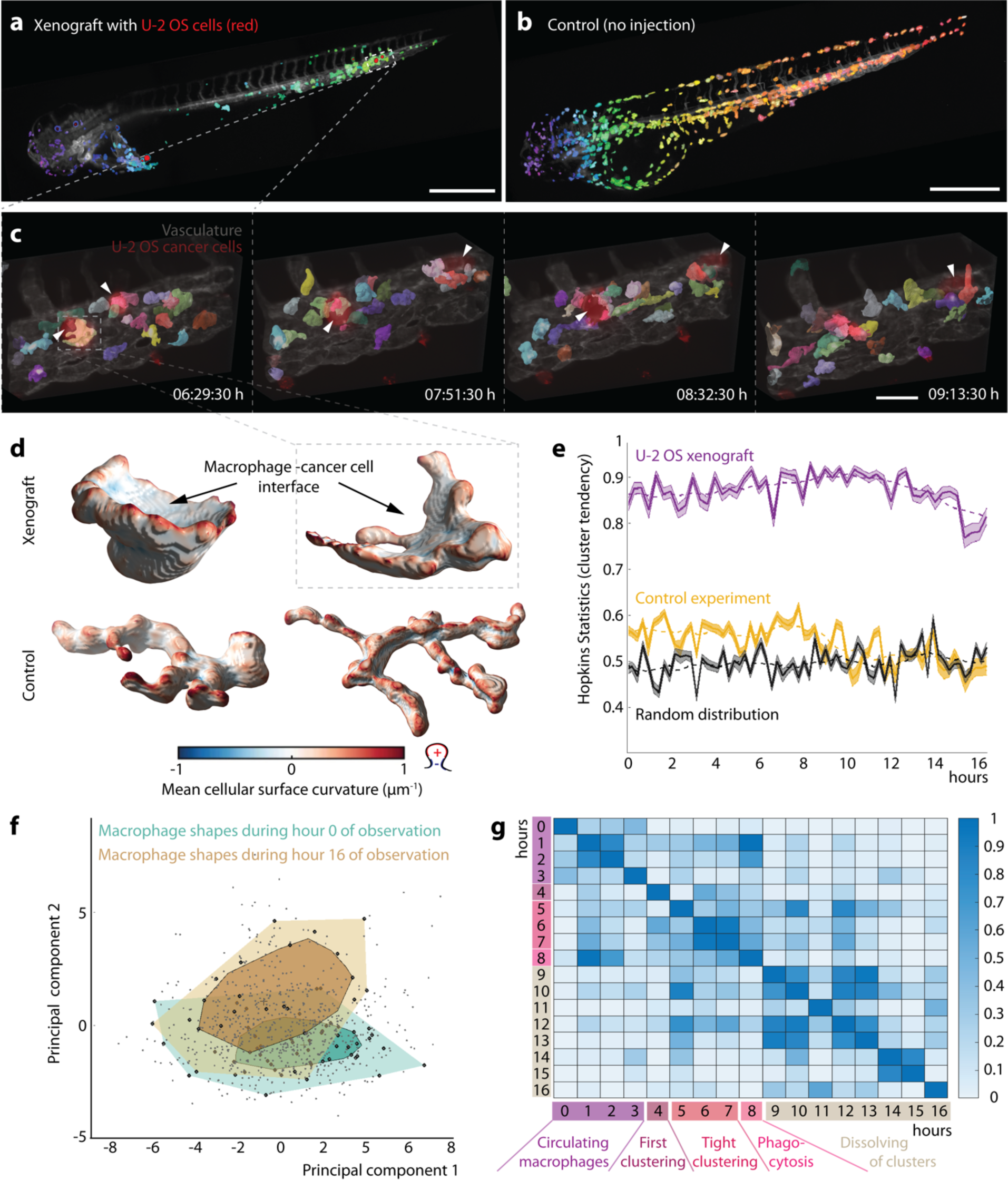
Analysis of self-driving, multi-scale data. The multi-scale data provided information on macrophage behavior from subcellular to organismal length scales in a U-2 OS osteosarcoma zebrafish xenograft (Supplementary Movie 4). **a** 3D rendering of low-resolution data of a zebrafish xenograft with U-2 OS osteosarcoma cancer cells (red) and segmented macrophages (color scale: violet to green from head to tail, Supplementary Movie 8). Macrophages clustered around sites with cancer cells. For tissue context, the vasculature (label: *Tg(kdrl:Hsa.HRAS-mCherry))* was rendered in gray. Boxed region indicates the location of high-resolution region in panel c. **b** 3D rendering of low-resolution data of a zebrafish embryo without xenograft (control, Supplementary Movie 9). All segmented macrophages distributed uniformly along the body axis (color scale: violet to red from head to tail). For tissue context, the vasculature (label: *Tg(kdrl:Hsa.HRAS-mCherry))* was rendered in gray. **c** 3D rendering of selected stills from high-resolution time-lapse data with segmented macrophages (in color, Supplementary Movie 10), U-2 OS cancer cells (red, white arrowhead) and vasculature (gray). Stills show concerted action of macrophages to phagocytose cancer cells. Boxed region indicates individual macrophage highlighted in panel d. **d** Representative 3D renderings of individual macrophages (Supplementary Movie 11) from the high-resolution data of xenograft experiment (top) and control (bottom). Color indicates mean cellular curvature. **e** Hopkins statistics to analyze macrophage dissemination inside a zebrafish with 10% sampling and dimensionality d=3, as a measure of cluster tendency, i.e. how clustered the macrophages were around cancer cells (violet). A higher number indicates a higher cluster tendency. Control (no xenograft) data is displayed in yellow. Simulation of a randomized, uniform distribution is shown in black. Data displayed as average ± 95% confidence interval (n=350 iterations for Hopkins statistic calculation). Dashed line indicates moving average of 10 timepoints. **f** Global morphological feature analysis of macrophage shapes from the high-resolution data of the U-2 OS cancer cell xenograft experiment over time, displayed in a principal component analysis (PCA) plot. Bagplots of all macrophage shapes in the first hour (hour 0, turquoise) and hour 16 (brown) of observation are overlaid for visualization and comparison. The bagplot consists of the Tukey median (cross), an inner polygon (darker color) that contains the 50% observations with largest Tukey depth, and the outer polygon (lighter color) with all data excluding outliers. Small gray points represent macrophage morphologies at all other timepoints. **g** Change of macrophage shapes over time: all macrophage shapes within an hour of observation were compared to macrophage shapes at different hours by applying a permutation test (n=300 iterations) using the Tukey median as a metric. The resulting p-values were plotted as a heat map. Scale-bar lengths are as follows: **a,b** 500 μm; **c** 50 μm.

To fulfill these requirements, our on-the-fly region tracking algorithm consisted of two steps, i.e., an initialization step at the start of imaging the high-resolution region of interest, and an update step from the previous timepoint to the current acquisition. During the initialization, we established correspondence using calibrated stage positions between the high-resolution region and the corresponding low-resolution data in any channel (fluorescence or brightfield). We found experimentally that increasing the tracked volume by 1.5 times in lateral directions compared to the size of the high-resolution region resulted in more robust tracking. To achieve fast tracking, we mapped the 3D volume of the tracked region into x-y, y-z, x-z maximum intensity projections and saved them in an image library. For each update, we applied multi-scale OpenCV’s template matching^21^ to find the previous x-y projected region in the current low-resolution data. Subsequently, the axial projections (y-z, x-z) were generated and registered to the axial projections of the previous timepoint. This resulted in the 3D shift vector of the region of interest and allowed us to update the stage positions for high-resolution imaging. Lastly, the image library was updated with the latest projections to account for changes in the region of interest over time.

### *In vitro* and *In vivo* imaging of development and disease processes

We demonstrate the application of our self-driving, multi-scale platform to a variety of biological processes in development and disease *in vitro* and *in vivo*. Specifically, we showcase an *in vitro* assay of cancer cell invasion using spheroid models, *in vivo* developmental processes of zebrafish gastrulation and vascular growth, and an *in vivo* larval zebrafish xenograft model to study processes of cancer cell metastasis. These biological processes extend over long-time periods and are inherently multi-scale in nature.

Invasion of a cancer cells from a primary tumor into the surrounding tissue is a critical stage of cancer progression^26^. Multi-scale imaging of this process is desirable to understand the overall behavior of a tumor, such as how many cells start migrating away from the tumor, while understanding the single cell processes that drive individual cells at the leading front. To study this process, cancer spheroids have become instrumental models^27^. We formed cancer spheroids in a Collagen matrix from SUM159 breast cancer cells that are known for their high invasiveness^28^. Using our self-driving, multi-scale imaging platform, we visualized the entire spheroid with low-resolution, and simultaneously imaged cancer cells (trailblazers) at the leading front with sub-cellular resolution. By mixing SUM159 cells of two colors 1:1 (Lifeact-GFP and the Lifeact-mCherry), individual cells were discernible. This allowed us to visualize actin reorganization during cell division at the leading front (Fig. 2a), while observing overall spheroid growth (Supplementary Movie 1).

In human development, starting from a single cell, a complex organism forms that consist of trillions of cells^46^. To shed light on this intricate process, multi-scale imaging is advantageous as tissue and organismal properties modulate the behavior of single cells. To study developmental processes, zebrafish are powerful and widely used model organisms^29^. They provide a short generation time, fast development and optical translucency, particularly in models without pigmentation^30^. This is in stark contrast to the highly scattering tissues in murine models, where fluorescence can only be superficially imaged with two or three-photon raster scanning microscopy^31^. To demonstrate the self-driving multi-scale microscope for imaging developmental processes, we visualized cellular divisions during zebrafish gastrulation (Supplementary Movie 2). While whole organism imaging allowed precise developmental staging and understanding of overall tissue flow, the high-resolution modality enabled precise characterizations of cell divisions, including sister chromatid separation (Fig. 2b). At a later developmental stage, we imaged the growing vascular network by combining high-resolution ASLM imaging of fluorescently tagged vasculature, labeled with *Tg(kdrl:EGFP),* with LED brightfield illumination for guidance. Over more than 11 hours, we followed the growth of the vasculature at the tip of the tail, while the whole embryo was growing and expanding over 100 μm. The resulting images revealed angiogenesis and anastomosis events, including rearrangements of the nuclei during this process (Extended Fig. 3).

Cancer metastasis is a major determinant of cancer-associated mortality and consists of a cascade of highly complex and dynamic processes during which cancer cells disseminate from their site of origin to invade distant organs ^32,33^. To study the dynamics and spatial context of these processes *in vivo*, multi-scale imaging is valuable to observe the dissemination and behavior of cells in different metastatic niches on a whole organism level, while imaging the sub-cellular processes that drive disease progression. To study cancer metastasis, zebrafish xenografts (Extended Fig. 4a) have been established as potent *in vivo* assays^34,35^. Zebrafish provide different metastatic niches and a physiological environment with endothelial cells, blood flow, and immune cells^36,37^. We leveraged this model to xenograft human breast cancer cells MDA-MB-231 that stably expressed a filamentous actin marker F-tractin-EGFP. For tissue context, zebrafish expressed the vascular marker *Tg(kdrl:Hsa.HRAS-mCherry)*. In the low-resolution mSPIM imaging, we visualized dissemination patterns, i.e. where cancer cells resided over time. The high-resolution ASLM imaging provided a detailed view on the cellular shapes, actin distribution and intricate network of protrusions that MDA-MB-231 cells formed inside the vasculature to attach to the vessel walls and other cells (Fig. 2c, Supplementary Movie 3).

### *In vivo* xenograft imaging to study macrophage-cancer cell interactions

To demonstrate the use of our self-driving, multi-scale microscopy platform for quantitative biology beyond visualization, we studied immune-cancer cell interactions in a zebrafish xenograft model. Zebrafish xenografts have been recently exploited to study interactions of cancer cells with cells of the innate immune system such as macrophages^38^. However, high-resolution images and quantification of these interactions at different scales over many hours are still lacking. We injected human U-2 OS osteosarcoma cancer cells, labelled with pVimentin-PsmOrange, into zebrafish larvae with fluorescent macrophages, labelled with *Tg(mpeg1:EGFP),* and *Tg(kdrl:Hsa.HRAS-mCherry)* labelled vasculature for tissue context (Extended Fig. 4). Two hours after injection, zebrafish with intact circulation and few isolated cancer colonies in the zebrafish tail were selected for imaging.

Our low-resolution, long-term time-lapse movie revealed overall macrophage dissemination in the xenografted larvae. Macrophages clustered around sites where cancer cells were present (Fig. 3a, Extended Fig. 4b, Supplementary Movie 4,5). In contrast, in zebrafish larvae without injected cancer cells, macrophages were uniformly distributed (Fig. 3b, Extended Fig. 4c, Supplementary Movie 6). Applying our self-driving microscope feature, the microscope observed and followed a selected micro-metastatic niche over 16 hours with high-resolution imaging to reveal how zebrafish macrophages interacted with and ultimately phagocytosed U-2 OS cancer cells, as the zebrafish larva grew (Fig. 3c, Extended Fig. 4d). Interestingly, several macrophages were present at the site of cancer cell ingestion, hinting at collective phagocytosis. While cancer cells formed a metastatic niche, they were remodeling the vascular network around them. After phagocytosis, the vasculature relaxed back to its normal state.

In contrast, in more highly metastatic cancer cells such as A375 melanoma cells, we did not observe such phagocytosis events over the same observation window despite close interactions between immune cells and A375 cells (Supplementary Fig. 9, Supplementary Movie 7). This was also reflected in the high survival rate of A375 cells in the tail over 48 hours after xenografting (Supplementary Fig. 10). However, we observed single apoptosis events of A375 cancer cells with subsequent clearance by macrophages (Supplementary Fig. 11; Supplementary Fig. 12).

To quantify the macrophage dynamics during phagocytosis of U-2 OS cancer cells across scales, we segmented the individual macrophages in both the low- and high-resolution data. To segment the low-resolution data, we used GPU-accelerated segmentation with py-clesperanto based on CLIJ^39^ (Fig 3a, b; Supplementary Fig. 13, Supplementary Movie 8, 9). However, the watershed-based approach of py-clesperanto failed to segment macrophages in the high-resolution data. This was due to the complex, non-spherical macrophage morphologies (Extended Figure 4d, Extended Figure 5a). Circulating and resident macrophages have large cytoplasmic branches, likely to survey as much tissue as possible for tissue damage or pathogens. Moreover, during phagocytosis, more compact macrophages clustered together (Figure 3c; Extended Figure 5a), rendering the separation of neighboring cells a difficult task. To overcome these challenges, we developed a custom segmentation workflow (Fig. 3c, Extended Figure 5c-g). This approach first used multi-Otsu thresholding and connected component labeling, to segment individual macrophages and identify macrophage clusters. To resolve individual macrophages inside the clusters, we complemented the initial segmentation with a 3D consensus segmentation based on 2D Cellpose^40^ segmentations of x-y, x-z, y-z views. Finally, manual curation was employed to resolve any segmentation ambiguities. The resulting segmentation provided a detailed characterization of macrophage morphologies in 3D over time (Supplementary Movie 10) and different conditions, e.g., xenograft vs control (Fig. 3d, Supplementary Movie 11). To further highlight the detailedness of the macrophage segmentation, we extracted a mesh of individual cell surfaces and calculated the mean cellular curvature on it (Fig. 3d).

The segmentations in low- and high-resolution data enabled a quantitative comparison of macrophage behavior across scales. In the low-resolution data, we quantified the spatial dissemination pattern of macrophages using the Hopkins statistic *H*^41^. Originally introduced to study plant distribution, the Hopkins statistic measures cluster tendency of the segmented macrophage positions against an equivalent number of randomly sampled positions in the zebrafish (Methods, Supplementary Fig. 14). A value of *H>*0.75, indicates at cluster tendency at the 90% confidence level^42^. The qualitative observation of clustering tendency of macrophages around cancer in the xenograft was confirmed by the high value of the Hopkins statistics over the 16 hours of time-lapse imaging. Except for one time point at the end of the recording, *H* was always above 0.75. In contrast, the macrophage dissemination without xenografted cancer cells was close to a uniform distribution (H< 0.6), and always less than in the xenograft experiment (Fig. 3e).

To systematically quantify the morphological shape changes of macrophages interacting with U-2 OS cancer cells, we used the segmentations of the high-resolution imaging data to perform a morphological feature analysis based on measurements of global cell shape features (Supplementary Table 2) ^36^. This analysis maps the macrophage shapes into a 2D principal components analysis (PCA) morphological shape space (Fig. 3f). Thereby, the first two components explain 82.7% of the observed variability in cell shapes, justifying the cut off after 2 components (Supplementary Fig. 15). In this 2D PCA space, the shapes of different macrophage populations could be compared with each other, for example by bagplots^43^, a generalization of univariate boxplots. To obtain insights into whether macrophages change their morphologies over the course of the time-lapse imaging, we grouped macrophage shapes defined by the hour of observation. Next, we compared each of these populations with each other using a permutation test based on the Tukey median as a metric^43^ (Fig. 3g), e.g., all macrophage shapes observed in hour zero vs. in hour sixteen. Interestingly, we observed that changes in macrophage shapes correlated with changes in biological function, i.e., circulating macrophages, first attachments, tight attachment, phagocytosis and dissolving of the clusters (Fig. 3g, Supplementary Movie 10). Taken together, our approach uncovers unique properties of cancer-cell interacting macrophages across spatial scales at biologically relevant temporal scales, demonstrating the utility and potential of self-driving multi-scale microscopy.

## Discussion

Development, cancer metastasis and immune-cancer cell interactions span multiple time and length scales^44–46^. To observe and quantify these biological processes from sub-cellular to organismal scales *in vivo* over long time periods, we have developed a self-driving, multi-scale light-sheet imaging platform. It enabled imaging of large 3D volumes over long time spans to reveal cell spreading and migration patterns, and to find rare dynamic events such as immune-cancer cell interactions. Guided by our own self-driving hardware control, the microscope could then selectively focus on processes of interest to concurrently visualize molecular signaling and morphological changes, such as cell shape changes, chromatid segregation, cell-cell interactions, or cytoskeletal rearrangements.

The need for multi-scale platforms arises as current microscopy platforms are limited in their spatiotemporal sampling due to technical and practical limitations^13^. Hence, available platforms do not allow the imaging of an entire living organism with sub-cellular resolution on relevant time scales. While smart microscopes have been introduced that combine different imaging modalities^15–17^, our platform is, to our knowledge, the first to allow for simultaneous imaging of complex 3D samples and organisms with concurrent, self-driving imaging of selected regions of interest with high-resolution. In our applications, we could select these regions of interest at the start of an experiment by careful experimental design and follow them over long time periods. For example, we observed and followed trailblazer cancer cells in cancer spheroids invasion assays, cells undergoing gastrulation, vascular cells in development, and cancer cells in metastatic niches in zebrafish xenograft models. In the future, we will extend the versatility of the platform by guiding the high-resolution data acquisition with event-based approaches.

An important aspect of our multi-scale platform is its self-driving feature, where the low-resolution imaging does not only provide biological information but also guides high-resolution imaging. Without this feature, high-resolution regions of interest quickly move out of view, which has hitherto prevented long-term, high-resolution acquisitions *in situ* in living organisms over more than few hours. In principle, instead of using low-resolution data for guidance, one could devise acquisition routines where the high-resolution region is imaged repeatedly as fast as possible and use motion prediction from high-resolution data alone to follow select region of interests. However, high-resolution imaging is considerably slower than low-resolution imaging, and typically requires a higher excitation duration and laser power. Therefore, for long-term imaging over many hours to days, phototoxicity and photobleaching stemming from high-resolution imaging needs to be minimized. Additionally, decoupling high-resolution imaging and guidance allowed for more options in experimental design, e.g., by combining bright-field transmission imaging of non-labelled large tissue structures for guidance (Extended Fig. 3). Importantly, our self-driving microscope control was process- and fluorophore-independent as it only relied on template matching to perform motion tracking, with no need for training data. This is distinct from deep learning methods for event-based imaging such as YOLO networks that need to be trained to recognize specific processes^17^. Our software, however, is designed to incorporate such machine learning algorithms in the future. Thereby, the availability of low-resolution data opens the path forward to search for specific processes of interest in an entire organism.

For the microscope user, the multi-resolution imaging capabilities provided an improved ease of use as regions of interest could be easily found within large volumes. This contrasts with dedicated high-resolution microscopes (in particular light-sheet architectures), which have a fixed, rather small field of view, which makes navigation within a 3D organism time-consuming and demanding.

As our system was designed for sub-cellular and organismal imaging *in vivo*, we leveraged a traditional light-sheet layout to optimize image performance. In turn, this constrained the sample mounting in comparison with recent open-top geometry implementations^12,47,48^ that may provide more flexibility in sample preparation. However, both the system design and sample holders can be adapted towards specific research questions. Because of the MVC design, our custom microscope hardware control is transferrable to other systems.

The presented data showcases applications of self-driving, multi-scale imaging across a wide range of biological questions, from developmental processes to processes in disease such as cancer. Specifically, we introduced self-driving, multi-scale imaging as a powerful platform to study immune cell – cancer cell interactions inside zebrafish xenograft models. In our experiments, we observed that U-2 OS cells were readily cleared by macrophages, while A375 evaded the immune system. Interestingly, these results mirror studies in murine models, where U-2 OS cells do not metastasize well ^49^, in contrast to A375 cells^50^. Thus, our zebrafish xenograft model provides an explanation of these observed metastasis patterns. Importantly, our platform enables follow-up studies of these interactions *in situ,* for the first time with the molecular detail necessary to understand what protects certain cancer cells from immune attack, while others are phagocytosed.

In summary, we have built, programmed, and applied a novel multi-scale microscope that enables observations of biological processes of interest from sub-cellular to organismal scales. We anticipate that our platform will find application and generate enriched datasets to address many biological questions in development and disease.

## Extended Figures

**Extended Figure 1.**
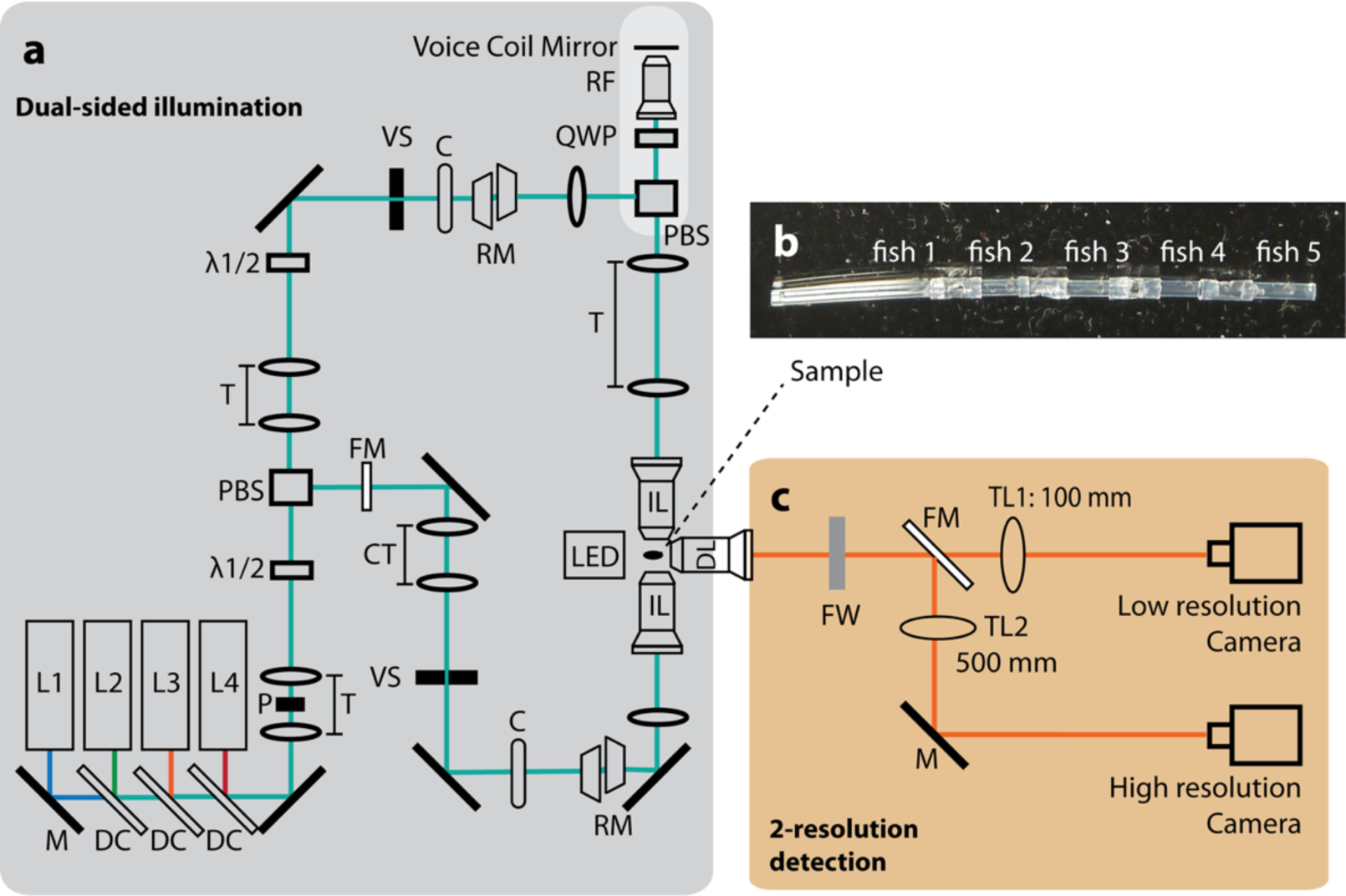
Schematic of the multi-scale microscope. **a** For the dual-sided illumination, first the laser lines of the 4 different lasers (L1-L4) were combined by dichroic beamsplitters (DC). Next, the beam was cleaned up with a pinhole (P) through which the light was focused with a 4x telescope (T). Then, a halfwave plate (λ 1/2) with a subsequent polarized beamsplitter cube (PBS) decided how much laser light went into which illumination arms. In the low-resolution illumination arm (bottom arm), a flip mirror (FM) could be switched into the illumination path to block the lower illumination during high-resolution imaging. Next, a telescope consisting of two cylindrical lenses expanded the beam in one direction to minimize light loss at the vertical slit (VS). A cylindrical lens formed the light-sheet that was pivoted by a resonant galvanometer (RM). Finally, another tube lens and an illumination objective (IL) formed the light sheet. In the high-resolution arm (top illumination arm), first the beam was further expanded by a telescope (T). Next, a halfwave plate (λ 1/2) polarized the beam so that all light was sent onto the refocusing unit (light gray). Before that, the beam was modified by a motorized vertical slit (VS), a light-sheet was formed with a cylindrical lens (C) and a resonant galvanometer (RM) generated a pivoting light-sheet. Next, another tube lens was conjugated with the remote focusing objective (RF). Thereby, the light was sent onto the voice coil mirror through a polarizing beamsplitter cube (PBS) and a Quarterwave plate (QWP). The remote focusing unit (light gray) was conjugated through another 4f system / telescope (T) to the illumination objective (IL) of the high-resolution arm. To position and image the sample in transmission (bright field), an LED illumination (LED) was available. **b** The microscope hardware comfortably accommodated several (zebrafish) samples in one experiment, mounted in low melting agarose with FEP tubes. **c** The detection was also composed of two arms. For the low-resolution detection, a 100 mm tube lens (TL1) was used. When changing to high-resolution imaging, a flip mirror (FM) sent the light through a 500 mm tube lens (TL2) onto the high-resolution imaging camera.

**Extended figure 2:**
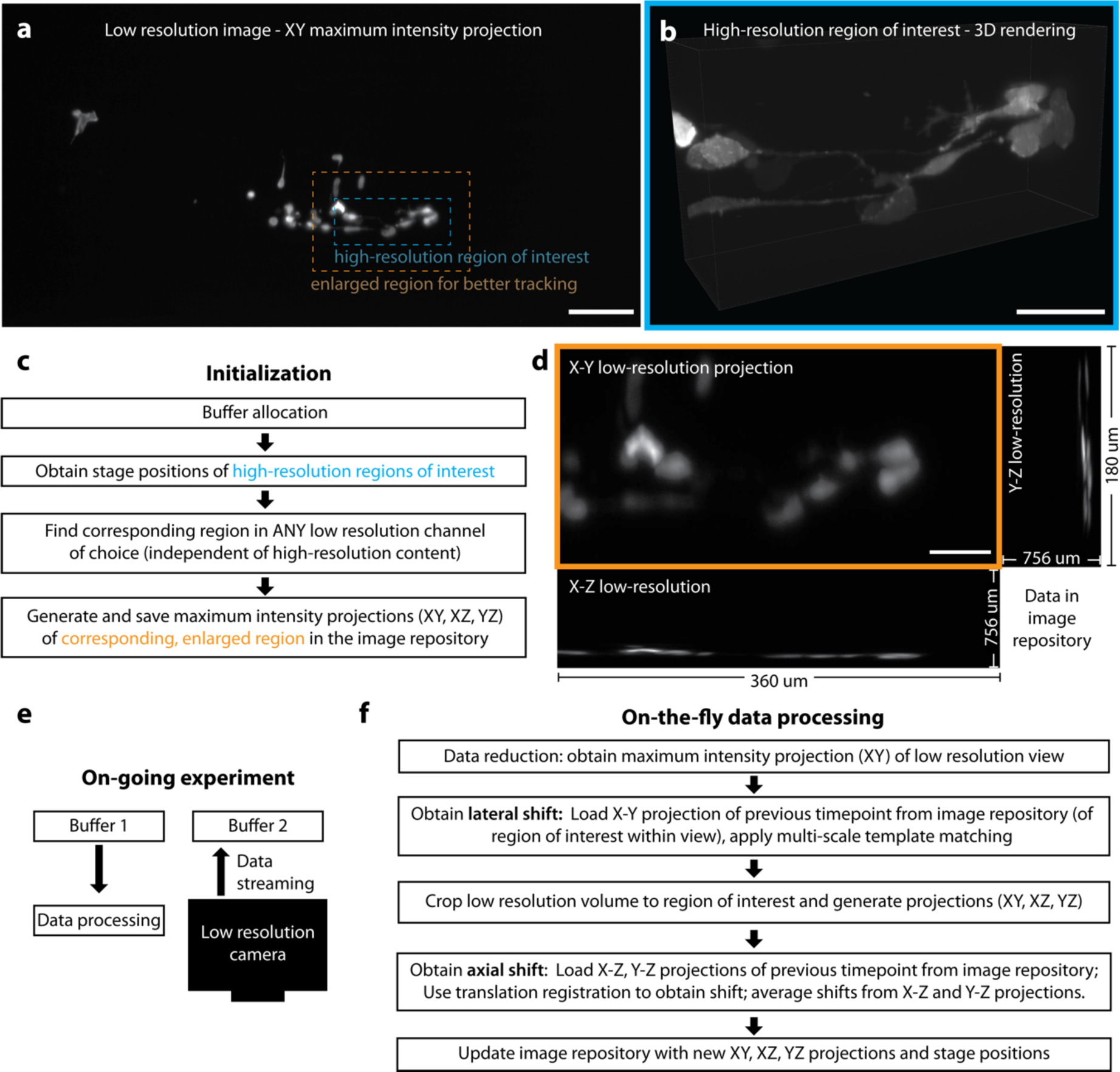
Region tracking algorithm. **a** Low-resolution maximum intensity projection of MDA-MB-231 breast cancer cells, labelled with F-tractin-GFP, in the zebrafish tail. Insets depict a high-resolution region of interest (blue, panel b) with its corresponding, enlarged (1.5x in both lateral dimensions) region used for better tracking. **b** 3D rendering of the high-resolution region of interest. **c** Schematic of the steps required for initialization of the region tracking algorithm. **d** X-Y, Y-Z, X-Z low-resolution maximum intensity projections saved in the image library at initialization, and each subsequent timepoint / update step. **e** Schematic of simultaneous data processing and acquisition (data streaming), enabled by a custom buffer architecture using shared memory arrays and tools from the concurrency tool library by Thayer, York *et al.*^22^**. f** Schematic of the pipeline for the update step to calculate on-the-fly the lateral and axial shifts for region tracking. Scale-bar lengths are as follows: **a** 150 μm, **b,d** 50 μm.

**Extended Figure 3:**
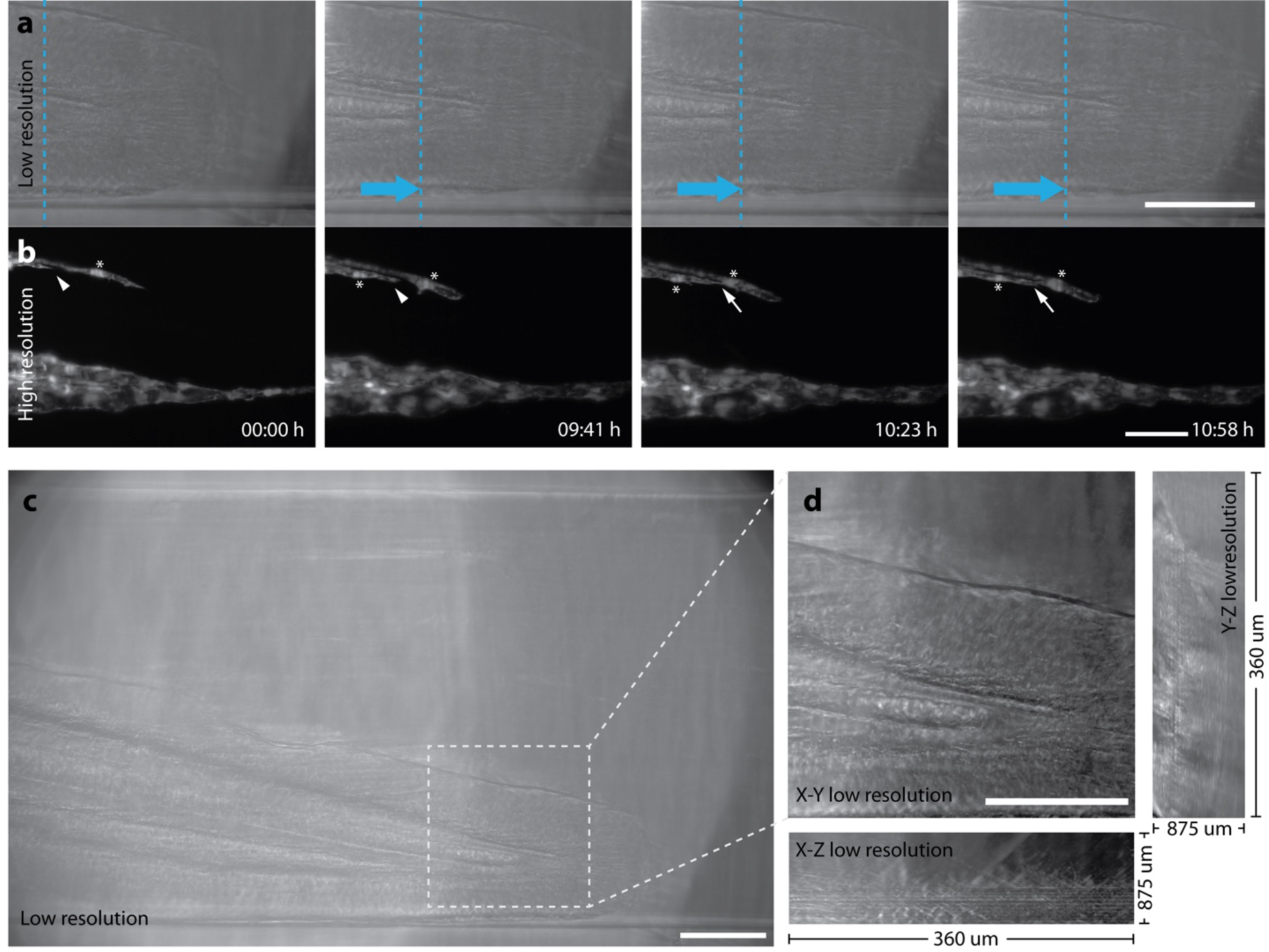
Region tracking using transmission images. **a** Selected maximum intensity projections of transmission images of the larval zebrafish tail over 11 hours of observation, starting at 2.5 days post fertilization. The blue dashed line indicates the tip of the vasculature at the end of the tail, growing around 100 μm over the 11 hours window of observation (blue arrows). **b** Despite this growth, self-driving microscopy kept the vascular region of interest in focus over the observation window. Maximum intensity projections of the corresponding high-resolution region showcase details of the outgrowth of a single vessel (white arrowhead) with subsequent anastomosis (white arrows) with a neighboring vessel, including positioning of endothelial nuclei (asterisk). The vasculature was labelled with *Tg(kdrl:EGFP)*. **c** Maximum intensity projection of the entire field of view of the low-resolution acquisition at start of the time-lapse imaging, with the inset highlighting the region used for tracking. **d** Representative X-Y, Y-Z, X-Z maximum intensity projections used in the region tracking algorithm. Scale-bar lengths are as follows: **a, c, d** 150 μm; **b** 50 μm.

**Extended Figure 4:**
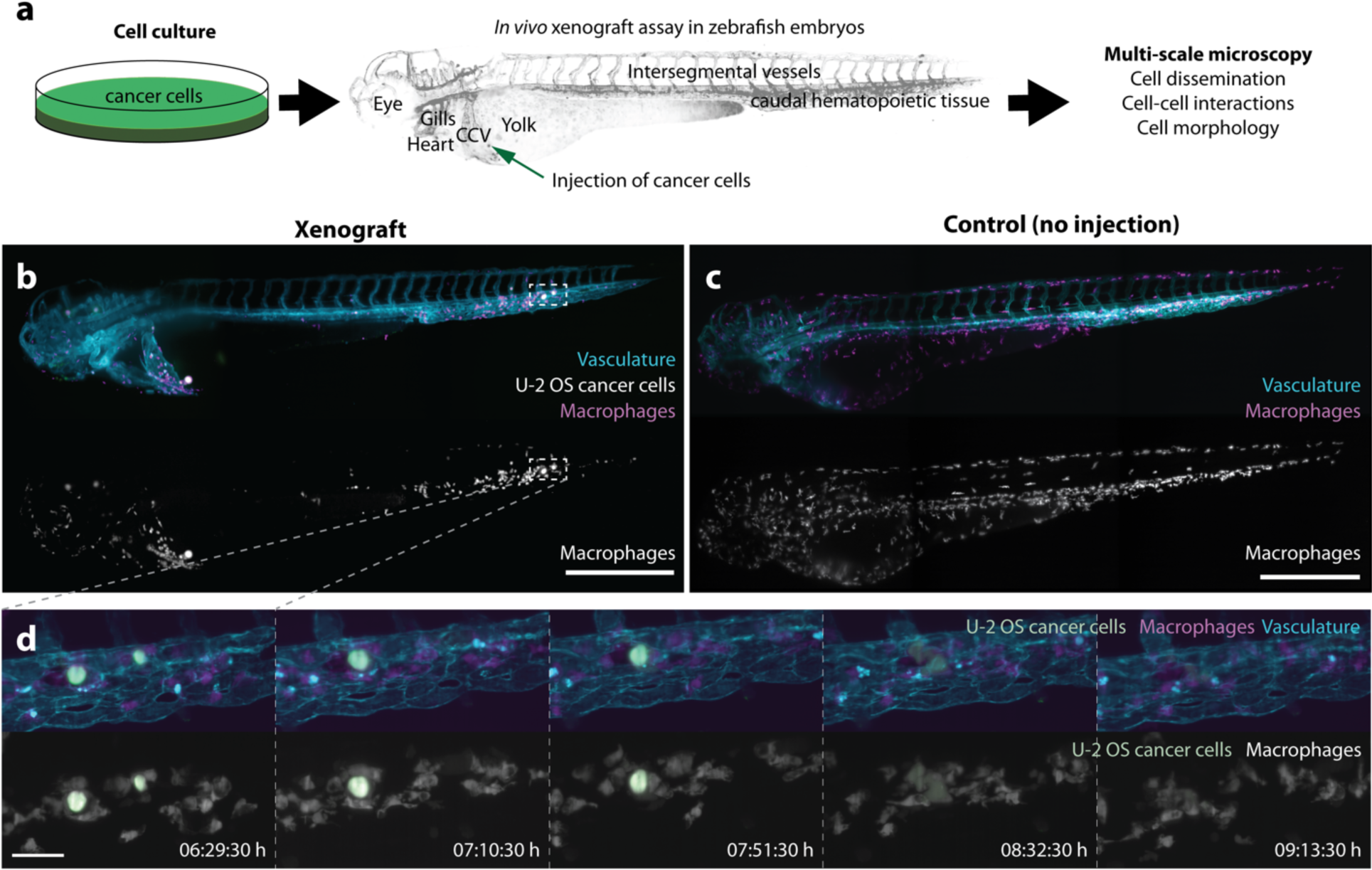
Phagocytosis of cancer cell by macrophages. **a** Schematic of our *in vivo* zebrafish xenograft assay to study immune cell–cancer cell interactions *in situ*. Firstly, human cancer cells were cultured and modified as desired, e.g., by expressing a fluorescent marker to label the cells. Secondly, cells were harvested and injected near the common cardinal vein (CCV) into the yolk of zebrafish larvae (violet arrow). Then, xenografts were imaged on our self-driving, multi-resolution microscope, and subsequent analysis allowed visualization and quantification of cell spreading, cell-cell interactions, and cell morphological changes. **b** Low-resolution mSPIM images captured the distribution of macrophages (*Tg(mpeg1:EGFP)*, top: magenta, bottom: grayscale image) in the entire zebrafish larvae after xenografting U-2 OS osteosarcoma cells (pVimentin-PsmOrange label, top: white, bottom: bright white). For tissue context, zebrafish also expressed the vascular marker *Tg(kdrl:Hsa.HRAS-mCherry)* (top: cyan). The image highlights how macrophages clustered around sites with cancer cells (Supplementary Movie 4). **C** In contrast, zebrafish without xenografts (control) displayed a uniform distribution of macrophages (*Tg(mpeg1:EGFP)*, top: magenta, bottom: grayscale image) across the entire zebrafish embryo (top: vascular marker *Tg(kdrl:Has.HRAS-mCherry* in cyan). **D** The self-driving feature of the microscope enabled high-resolution imaging of selected cancer colonies in the zebrafish tail over many hours by keeping it in focus. Frequently, we observed how zebrafish macrophages (*Tg(mpeg1:EGFP)*, top: magenta, bottom: gray) attached to the U-2 OS cancer cells (pVimentin-PsmOrange label, top and bottom: green), and phagocytosed them (Supplementary Movie 4, 5). Scale-bar lengths are as follows: **b,c** 500 μm; **d** 50 μm.

**Extended Figure 5:**
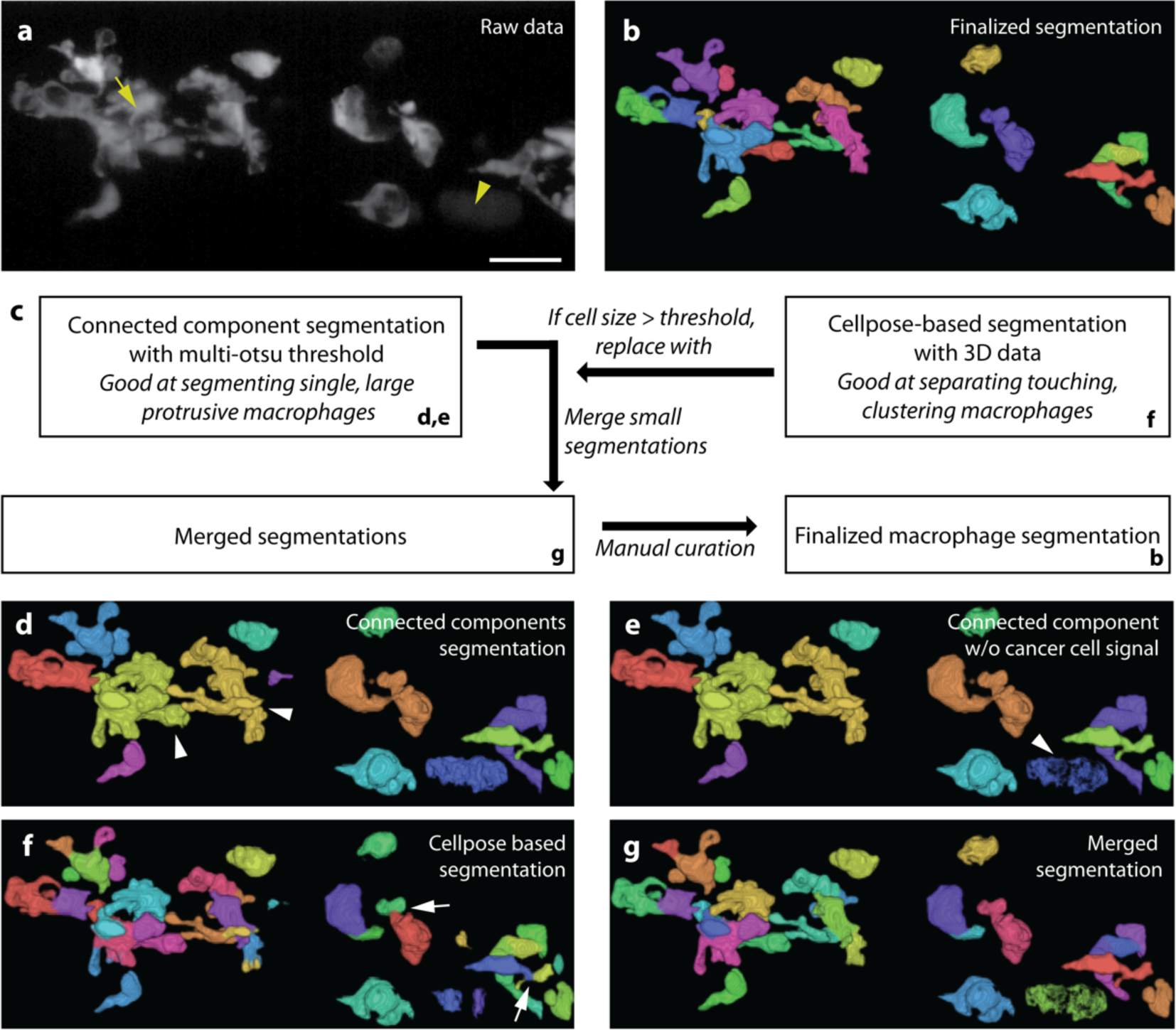
Workflow to obtain a high-resolution segmentation of zebrafish macrophages. **a** Maximum intensity projection of a 3D volume in the caudal hematopoietic tissue visualizing zebrafish macrophages, labeled with *Tg(mpeg1:EGFP)*^51^, after xenografting human U-2 OS cells into the zebrafish larvae. As the cancer cells (yellow arrowhead) were labelled with a psmOrange fluorophore, there was bleed-through into the macrophage channel (GFP). Moreover, macrophages tended to cluster around cancer cells and at locations where cancer cells resided before they were phagocytosed by macrophages (yellow arrow). **b** 3D rendering with Fiji’s 3D viewer to display the final segmentation by the proposed workflow. **c** Schematic of the segmentation workflow. **d** We initially segmented the data with multi-Otsu thresholding and connected component labeling. This segmented individual macrophages well, e.g. in the control experiment (without xenografts) or at the beginning of the xenograft experiment. However, it could not separate touching macrophages (white arrowheads). **e** To clean up the segmentation, we first removed the cancer cell signal (white arrowhead) from the segmentation. **f** To separate clustering macrophages, we constructed 3D consensus segmentation from Cellpose-2D^40^ segmentations computed in orthogonal x-y, x-z, y-z views of the original raw data. While this segmentation separated clustering macrophages, it was observed to split individual macrophages into smaller fragments (white arrows) when the macrophages show extensive cytoplasmic extensions of several tens to hundreds of microns. **g** We merged the connected component segmentation and the Cellpose-based segmentation by replacing connected volumes that exceeded a defined threshold with the Cellpose-based segmentation. Moreover, we merged connected volumes that were below a certain threshold with neighboring volumes. Subsequent manual curation further cleaned up the segmentation of macrophages to obtain the finalized segmentation **b**. Scale-bar lengths are as follows: **a** 30 μm.

## Methods & Protocols

The research within this work complies with all relevant ethical regulations as reviewed and approved by the University of Texas Southwestern Medical Center.

### Microscope layout

A schematic of the microscope is available as Extended Fig. 1 with a detailed list of all microscope hardware components available in Supplementary Table 1. Moreover, both illumination light paths are characterized in more detail in Supplementary Figures 1 and 2. Acquired data was stored on the BioHPC cluster, provided by UT Southwestern Medical School.

### Microscope control software

The microscope was controlled by a custom Python-based microscope control software based on an MVC design pattern (Supplementary Fig. 7). It featured concurrent acquisition and processing, an intuitive graphical user interface with a state-of-the art image viewer for ease of use (Supplementary Fig. 8), automated generation of maximum intensity projections for fast browsing of the acquired data, and control of different light-sheet modalities including mSPIM and ASLM. The control code is available as Github repository.

### Fluorescent nanosphere measurements

Fluorescent nanosphere samples of 0.2 *μ*m YG Nanospheres (Polyscience, 17151) were prepared in 2% low melting agarose (Sigma Aldrich, A9045) at a concentration of 1:1000 – 1:10’000. After imaging, the point spread function (Fig. 1c, Supplementary Fig. 3) was determined using the plugin MetroloJ^52^ in Fiji^53^ (n=20 for each resolution level). The mean and confidence interval (95%) were calculated using R, and the data was checked for normality of the measurements (Supplementary Note 2).

### Zebrafish husbandry and sample preparation

Zebrafish husbandry and experiments described here have been approved and conducted under the oversight of the Institutional Animal Care and Use Committee (IACUC) at UT Southwestern under protocol number 101805. Zebrafish (*Danio rerio*) adults and embryos were kept at 28.5°C and were handled according to established protocols^54,55^. All zebrafish experiments were performed at the embryonic stage and therefore the sex of the organism was not yet determined.

To visualize the development of the zebrafish vasculature, the zebrafish line expressing the vascular marker *Tg(kdrl:EGFP)*^56^ was imaged. To visualize macrophage interaction with cancer cells, the fish line *Tg(mpeg1:EGFP)*^51^ was crossed with zebrafish expressing the vascular marker *Tg(kdrl:Hsa.HRAS-mCherry)*^57^ in a casper background^30^. To visualize the nuclei inside developing zebrafish, the fish line *Tg(h2afva:h2afva-GFP)*^58^ was used.

To immobilize the zebrafish for imaging, zebrafish embryos were anesthetized with 200 mg/l Tricaine (Sigma Aldrich, E10521)^59^ during imaging. To mechanically mount up to five zebrafish embryos for imaging, they were individually embedded in 0.1% low melting agarose (Sigma Aldrich, A9414) within fluorinated ethylene propylene (FEP) tubes (Pro Liquid GmbH, Art: 2001048_E; inner diameter 0.8 mm and outer diameter 1.2 mm), coated with 3% methyl cellulose (Sigma Aldrich, M0387), and connected as previously described^10^.

### Spheroid sample preparation

SUM159 breast cancer cells^27,60^ were cultured in DMEM/F12 media (Gibco, 11320033) containing 5% FBS (Peak Serum), 1ug/ml hydrocortisone (Sigma Aldrich, H0888), 5ug/ml bovine insulin (Sigma Aldrich, I6634), and 1% Penicillin-Streptomycin (Gibco, 15140122). A Lifeact-GFP and a Lifeact-mCherry expressing SUM159 cell line were established through lipofectamine transfection according to the manufacturer’s protocol. pTK92_Lifeact-GFP (Addgene plasmid # 46356) and pTK93_Lifeact-mCherry (Addgene plasmid # 46357) were obtained from Addgene.

To obtain mosaic-labelled cancer spheroids, a 1:1 mixture of the Lifeact-GFP and the Lifeact-mCherry SUM159 cell lines (1000 cells each) was incubated in 96-well Nunclon Sphera low adhesion plates (Thermo Fisher Scientific, 174925) for 48 hours at 37°C and 5% CO_2_. The spheroids were then resuspended in a 50 uL mixture of 70% activated rat tail Collagen 1 (Corning, catalog no. 354236) and 30% Cultrex UltiMatrix Basement Membrane Extract (R&D systems, BME001) at a final concentration of 2.1 mg/ml Collagen 1 and 3mg/ml UltiMatrix BME. The resuspended spheroids were then embedded in sterile FEP tubes containing holes punctured with 30 G needles for increased circulation. FEP sterilization was achieved by flushing the tubes with 70% ethanol and letting them dry in a closed plate in the incubator. After 24 hours of additional incubation at 37°C and 5% CO_2_, the spheroids were imaged.

### Cell culture for zebrafish xenograft

A375 cells^61^ (ATCC CRL1619) and MDA-MB-231^63^ cells (ATCC HTB-26) were acquired from ATCC. U-2 OS cells^62^ were obtained from Richard McIntosh (University of Colorado, Boulder CO). To test for mycoplasma contamination, the test kit Genlantis MycoScope PCR Detection Kit (MY01100) was used. U-2 OS, MDA-MB-231, and A375 cells were cultured in DMEM-high glucose (Sigma Aldrich, D6429) supplemented with 10% fetal bovine serum (FBS; Peak Serum) and 1% antibiotic-antimycotic (Gibco, 15240062) at 37 °C and 5% CO_2_.

To complement the zebrafish macrophage marker *Tg(mpeg1:EGFP)*^51^ and the vascular marker *Tg(kdrl:Hsa.HRAS-mCherry)*^57^, we expressed the fluorescent marker pVimentin-PSmOrange^64^ in the xenografted U-2 OS and A375 cancer cells. pVimentin-PSmOrange was obtained from Addgene (plasmid # 31922). The MDA-MB-231 cells stably expressed F-tractin-EGFP^65^ to label the cells and reveal their actin organization. The F-tractin-EGFP construct was obtained from Addgene (plasmid # 58473).

To obtain fluorescently labelled cancer cells, U-2 OS cells were transfected with Lipofectamine LTX&PLUS Reagent (Invitrogen, 15338100) according to manufacturer’s protocol. On day 0, 300’000 cells were seeded per well of a 6-well plate (ThermoFisher, 140675). On day 1, cells were transfected in Opti-MEM Reduced Serum Medium (Gibco, 31985062) with Lipofectamine LTX&PLUS Reagent (Invitrogen, 15338100) at a concentration 3.5 ug DNA, 3.5 ul Plus and 12 ul LTX reagent. On day 2, the media was changed back to normal cell culture media and cells were selected using antibiotic resistance to neomycin (Gibco 10131035). A375 cells were transfected on day 1 with FuGENE HD (Promega, E2311) according to the manufacturer’s protocol. 150ul/well transfection mixture was prepared with 3 ug DNA and 9 ul FuGENE® HD Transfection Reagent in sterile water and 10 min complexing time at room temperature.

### Zebrafish xenografts

To prepare for a successful xenograft and imaging experiment, zebrafish embryos were collected and treated with 0.1 mM N-Phenylthiourea (PTU, Sigma Aldrich P7629) starting at 1 day post fertilization (dpf) to prevent pigmentation. At 2.25 dpf, the dechorionated embryos were xenografted with cancer cells. For this, cells were harvested at 70-90% confluency and trypsinized for 3 min (Gibco, 15400-054). If cells were highly adherent, cells were run through a 70-µm cell strainer (Fisherbrand, 22-363-548). To achieve an optimal number and density of cells for microinjection, 4 × 10^6^ cells in 40 uL normal cell culture media were prepared and stored on ice until xenografting.

Xenografts were generated by injection of the prepared cell mixture into the yolk near the common cardinal vein (CCV) zebrafish larvae with glass capillary needles (World Precision Instruments, 1B100-4), pulled on a micropipette puller (Sutter Instrument, P-1000). Thereby, 50-1000 cells were injected per fish. During the injection, zebrafish were anesthetized with Tricaine (Sigma Aldrich, E10521). Two hours after injection, zebrafish embryos with few single cancer cells in the zebrafish tail and intact circulation were selected under a fluorescent stereomicroscope for imaging.

### Low-resolution image processing and segmentation of zebrafish xenografts

Acquisition of three individual low-resolution tiles was required to cover the entire zebrafish larva. To fuse the three individual tiles to one 3D volume, we developed our own Python based stitching workflow. Firstly, we determined a coarse positioning of the tiles using the physical coordinates of the microscope stage position. Secondly, we calculated the maximum intensity projections of each tile to reduce the computational workload. Next, we calculated Fast Fourier transform-based cross-correlations on the overlapping sections of the background subtracted maximum intensity projections and obtained the translational shift between tiles as the highest value of all cross-correlations. Lastly, we fused the tiles using a sigmoidal blending function.

Next, we segmented the fused images to obtain the positions and dissemination patterns of macrophages (Supplementary Fig. 13). To segment the low resolution macrophage images, we used GPU accelerated segmentation with py-clesperanto based on CLIJ^39^ (https://github.com/clEsperanto/pyclesperanto_prototype). Segmentation parameters for the low-resolution segmentation were tuned with a graphical user interface, Napari (https://github.com/haesleinhuepf/napari-workflow-optimizer).

Specifically, before applying the segmentation workflow, we rescaled the data to isotropic voxel spacings to obtain accurate segmentations. Thereby, we downsampled the lateral pixel spacings to fit the larger axial plane-to-plane spacings. This allowed us to compress the data from ∼7 GB to ∼85 MB, enabling fast GPU accelerated processing on the local BioHPC cluster computer, equipped with Tesla V100 32GB GPU cards.

The segmentation workflow comprised pre-processing, thresholding, labeling with Voronoi-Otsu labeling on the binary image, and post-processing. To pre-process the data, we applied a top-hat filter for background subtraction with radius 6 in all directions. We then thresholded the image with a manually optimized threshold in Napari, and then applied Voronoi-Otsu labeling, which included two Gaussian blurs, spot detection, Otsu-thresholding and Voronoi labeling. Thereby, we used 2.0 as spot sigma and 1.0 as outline sigma as parameters in the Voronoi-Otsu labeling. Next, we removed spurious labels below a size of 10 voxels. Lastly, we saved the segmentation statistics and outcome parameters as an xlsx file for further analysis. The workflow can be found in the following Python file on Github: *segment_lowres_3d_stitched.py*.

### Hopkins statistics for analysis of spatial distribution

To quantify the cluster tendency of macrophages, we obtained the centroid position of each cell in the 3D volume from the low-resolution macrophage segmentations (real positions). Moreover, we calculated the 3D zebrafish volume *V* as a space for possible locations of macrophages to compare the observed spatial distribution (real positions) against randomly generated distributions (random positions) (Supplementary Fig. 14). We used these data points to test for evidence that the observed macrophage distributions contained clusters. To this end, we computed the Hopkins statistics *H*^41^, defined as^66,67^:

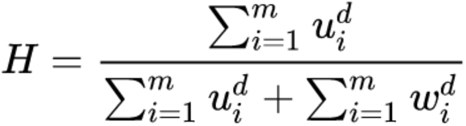

where *u_i_* is the Euclidean distance from a random position within the shape of the zebrafish *V* to the nearest-neighbor location of a real segmented macrophage position, and *w_i_* the Euclidean distance from a randomly selected real segmented macrophage position to the nearest-neighbor location of a real segmented macrophage position. The dimensionality d was set to three for the three spatial degrees of freedom associated with the macrophage positions. The number of random samples m calculated in the Hopkins statistic H was set to 10% of the total number of macrophages *n* for each time point. To account for variability in the random sampling, we calculated the Hopkins statistics 350 times, which we found to be a stable number of iterations, and determined the 95% confidence interval for display.

### High-resolution image segmentation of zebrafish xenograft data

To analyze macrophage morphology from the high-resolution data (Fig. 3d, Extended Fig. 5a), a multi-step segmentation pipeline was required (Extended Fig. 5b, c). First, we segmented the high-resolution data using multi-otsu thresholding^68^ from the scikit-image library^69^, with subsequent connected component labeling (Github: *ConnectedComponent_MacrophageSegmentation.py*). To optimize the segmentation, we performed data preprocessing, including an optional registration of the raw dataset using PyStackReg^70^, background smoothing, normalization, and Wiener-Hunt deconvolution with scikit-image. This segmentation performed well at identifying single, protrusive macrophages. Next, as the psmOrange fluorophore labelling cancer cells showed bleed-through to the macrophage channel (488 nm excitation), we removed the cancer cell signal from the images (*postprocessing_removeCancersignal.py*).

However, this approach did not distinguish individual macrophages that touch, e.g. during phagocytosis (Extended Fig. 5d). Therefore, we complemented the multi-otsu thresholding with deep learning based segmentation. For this, we used Cellpose^40^ and its cytoplasm 2.0 model (“cyto2”) to compute segmentations slice-by-slice in x-y, x-z, y-z views, and aggregated the resulting 2D segmentation into a single consensus 3D segmentation using custom Python code (*Cellpose_based_MacrophageSegmentation.py*). The full mathematical principles of this segmentation framework, suitable for a wide range of 3D cell imaging data along with in-depth validation and determination of method applicability will be described elsewhere.

To merge both segmentations (*Merge_ConnectedComponents_Cellpose.py*), we identified macrophage segmentation volumes in the multi-otsu thresholding approach that exceeded a defined threshold in size (40’000 voxels). The threshold was manually chosen based on a histogram distribution of the macrophage size distribution in the control data (i.e. no cancer cell xenograft), where macrophages rarely interacted with one another. We then replaced the label of the multi-otsu thresholding approach with the labels of the Cellpose-based segmentation. After replacement, we postprocessed the merged segmentation by merging labels below a defined threshold (5000 voxels) with the closest label.

Lastly, as macrophage segmentation was very challenging due to their clustering and shape differences, we performed manual curation of the dataset (*Manual_curation.py*) with a Python based script and Fiji^53^.

### Analysis of macrophage morphology over time

To analyze the morphological variation of macrophages over time, we performed global morphological feature analysis in Matlab vR2023b (MathWorks, Inc.), as previously described by Segal *et al.*^36^. We included 12 geometric features, including volume, surface area, solidity, sphericity, longest length, extend, aspect ratio, roughness, volume sphericity, radius sphericity, ratio sphericity, and circumscribed sphere area ratio (Supplementary Table 2). To visualize and analyze the resulting data in a 2D morphological shape space, we reduced the dimensionality of the features using principal component analysis (PCA).

To assess the shape variation of macrophages over time, we grouped data from one hour of imaging (three time points), and compared the spatial distribution of each group across all time points in the 2D principle component analysis (PCA) space. To assess the similarity between each pair, a permutation test with 300 iterations was employed. This involved the random shuffling of the positions in the 2D PCA space, and the subsequent measurements of the Euclidean distance between the Tukey median of each new distribution (see Rouseeuw et al.^43^). Similar distributions have a zero Tukey median difference.

### Analysis and rendering of macrophage curvature

The high-resolution data provided enough resolution to analyze the curvature of the macrophage cell surface. For this, the surface mesh was extracted from the 3D image segmentation using marching cubes with the ‘isosurface’ function in Matlab vR2023b (MathWorks, Inc.). The resulting mesh was smoothed with the Taubin filter^71^ using the ‘smoothSurfaceMesh’ function in MATLAB and the mean curvature was computed using principal curvatures as described by Rusinkiewicz^72,73^. For visualization, we used u-shape3D^74^ to color the triangle mesh using a blue, white, red colormap corresponding to negative, flat and positive curvature for a mean curvature range of [-1µm^-1^ , 1 µm^-1^]. The mesh was rendered using ChimeraX v.1.2.5^75^.

### Image and movie visualization

To visualize the high- and low-resolution data, we leveraged different software packages. For display, maximum intensity projections and 2D images were contrast, level and gamma adjusted in Fiji/ ImageJ^53^ and Adobe Photoshop. Optionally, a Gaussian blur and/or sharpen operation in Fiji was applied. 3D renderings were generated with Fiji’s 3D Viewer, Agave Renderer (https://github.com/allen-cell-animated/agave) and Napari^23^. Movies were annotated with boxes and arrows using a dedicated Fiji plugin^76^.

## Data availability

Due to the large size of the imaging datasets collected within this manuscript, the datasets are only available from the authors upon request.

## Code availability

All algorithms, code and software used in this study are available on Github. The microscope control code and all image processing software is available at: https://github.com/DaetwylerStephan/selfdriving-multiscale-imaging

## Acknowledgments

We would like to thank Andrew York for discussion on the control software, Bo-Jui Chang, Justine Keth, Dana Kim Reed and Kushal Bhatt for support with reagents, sample preparations, and sorting, the Animal Resource Center (ARC) for taking care of the zebrafish facility, and the whole Fiolka, Dean, Amatruda and Danuser labs for feedback and comments. Moreover, this research was supported in part by the computational resources provided by the BioHPC initiative at UT Southwestern Medical Center.

Funding for this work is acknowledged as follows: Swiss National Science Foundation, grant number 191347 to S.D.; National Institute of General Medical Sciences, grant number R35 GM133522 to R.F.; National Cancer Institute, grant number U54 CA268072 to G.D. and R.F; and grant number K99CA270285 to D.S.

## Ethics declarations

The authors declare no competing interests.

## Contributions

SD and RF wrote the manuscript and built the microscope, SD programmed and applied the microscope, SD, HMF and FZ performed data analysis, SD and ES performed sample preparations, JMW and RAB prepared cancer spheroids, DS and GD contributed to study design, SD, RF and GD secured funding for the project, all authors revised and approved the manuscript.

