## Supplementary Document for "Imaging of cellular dynamics *in vitro* and *in situ*: from a whole organism to sub-cellular imaging with self-driving, multi-scale microscopy"

### Overview

Supplementary Movies 1-12
Supplementary Note 1: Microscope Hardware
Supplementary Note 3: Fluorescent Nanosphere Measurements
Supplementary Note 2: Microscope Control Software Architecture
Supplementary Note 4: A375 Xenograft Experiments
Supplementary Note 5: Data Analysis
Supplementary Tables 1-2
Supplementary Figures 1-15
Supplementary References

### Supplementary Movies

**Supplementary Movie 1:** Multi-scale, time-lapse imaging of a mosaic-labelled SUM 159 breast cancer cell spheroid embedded into a collagen matrix with the low-resolution modality (top) and the high-resolution modality (bottom). The spheroids consisted of a 1:1 mixture of cells expressing the actin marker Lifeact-GFP (cyan) and Lifeact-mCherry (magenta). Boxed region on top indicates the location of the high-resolution region on the bottom. The yellow arrowhead points at a cell division at the invasive front. Maximum intensity projections of the timepoints are shown. Scale-bars: 150 µm (top), 20 µm (bottom).

**Supplementary Movie 2:** Multi-scale, time-lapse imaging of zebrafish gastrulation with cells expressing the histone marker *Tg(h2afva:h2afva-GFP),* starting at around 6 hours post fertilization. Low-resolution imaging (top) captured the entire zebrafish embryo (color scale: depth of data in 3D volume from 0 to 450 um), while the high-resolution imaging (bottom, maximum intensity projection) enabled near-simultaneous imaging of cell division. Scale-bars: 250 µm (top), 25 µm (bottom).

**Supplementary Movie 3:** Multi-scale, time-lapse imaging of human breast cancer cells MDA-MB-231 expressing F-tractin-EGFP (top: magenta, bottom: gray) *in situ* in a larval zebrafish xenograft model, expressing the vascular marker *Tg(kdrl:Hsa.HRAS-mCherry)* for tissue context (top: cyan). Low-resolution imaging (top, maximum intensity projections) captured the whole organism, while high-resolution imaging (bottom, maximum intensity projections) revealed sub-cellular dynamics including an intricate network of protrusions. The cancer cells were xenografted into zebrafish larvae at 2.25 days post fertilization. Scale-bars: 250 µm (top), 25 µm (bottom).

**Supplementary Movie 4:** Multi-scale, time-lapse imaging of U-2 OS osteosarcoma cancer cells (pVimentin-PsmOrange label, top: white, bottom: green) and their interactions with macrophages (*Tg(mpeg1:EGFP)*, magenta) in a larval zebrafish xenograft model. The zebrafish expressed the vascular label *Tg(kdrl:Hsa.HRAS-mCherry)* (cyan) for tissue context. Maximum intensity projection of the low-resolution data is shown on top, while a 3D rendering of the high-resolution is shown on the bottom. Boxed region on top indicates the location of the high-resolution region on the bottom. Scale-bars: 500 µm (top), 50 µm (bottom).

**Supplementary Movie 5:** High-resolution time-lapse movie of U-2 OS osteosarcoma cancer cells (pVimentin-PsmOrange label, top and bottom: green) xenografted into larval zebrafish with labelled macrophages (*Tg(mpeg1:EGFP),* top: magenta, bottom: gray) and vasculature (*Tg(kdrl:Hsa.HRAS-mCherry),* top: cyan), displayed as maximum intensity projections over time. Same high-resolution data as in Supplementary Movie 4. Scale-bar length: 30 µm.

**Supplementary Movie 6:** Low-resolution, time-lapse imaging of macrophages (*Tg(mpeg1:EGFP)*, magenta) in larval zebrafish expressing the vascular marker *Tg(kdrl:Hsa.HRAS-mCherry* (cyan) without any xenografted cancer cells (control). Scale-bar: 500 µm.

**Supplementary Movie 7:** High-resolution time-lapse movie of A375 melanoma cancer cells (green, pVimentin-PsmOrange label) and their interactions with macrophages (*Tg(mpeg1:EGFP)*, gray) (Supplementary Figure 9), displayed as 3D rendering over time. Scale-bar: 50 µm.

**Supplementary Movie 8:** 3D rendering of the segmented macrophages (color scale: violet to green from head to tail) from the low-resolution imaging data of the xenograft experiment with injected U-2 OS osteosarcoma cancer cells (red) (Supplementary Movie 4). For tissue context, the vasculature (label: *Tg(kdrl:Hsa.HRAS-mCherry))* was rendered in gray. Scale-bar: 500 µm.

**Supplementary Movie 9:** 3D rendering of the segmented macrophages (color scale: violet to red from head to tail) from the low-resolution imaging data over time in the control experiment (i.e. no injected cancer cells, Supplementary Movie 6). For tissue context, the vasculature (label: *Tg(kdrl:Hsa.HRAS-mCherry))* was rendered in gray. Scale-bar: 500 µm.

**Supplementary Movie 10:** Left: 3D rendering of the segmented macrophages (in color) from the high-resolution imaging data of the U-2 OS osteosarcoma cancer cell (in red) xenograft experiment (Supplementary Movie 4) with vasculature rendered in gray. Right: Analysis of macrophage shapes using global morphological feature analysis, displayed in the principal component (PCA) space. Each gray point represents a single macrophage. Bagplots were used to visualize all macrophage shapes within an hour of observation in the PCA space (Tukey median (cross); inner polygon (darker color) that contains the 50% observations with largest Tukey depth; the outer polygon (lighter color) with all data excluding outliers). The color of the bagplots is representing the classification of macrophage function from Fig. 3g. The red line indicates the change of the Tukey median over time. Scale bar: 50 µm

**Supplementary Movie 11:** Representative 3D renderings (not to scale) of individual macrophages from xenograft (top) and control (bottom). Color indicates mean cellular curvature.

**Supplementary Note 1: Microscope Hardware and performance**

In this note, we detail the hardware components, the beam paths, and camera configuration. In **Supplementary Table1**, a complete parts list of the optical and opto-electronic components of the multi-scale microscope are given.

**Supplementary Figures 1-2** show the excitation beam path for the low and high-resolution mode, respectively. The focal length of each lens and the beam size is indicated. In the low-resolution mode, shown in **Supplementary Figure 1**, the input laser beam was first expanded four-fold. A subsequent cylindrical lens pair expanded the beam in one dimension by another factor of four. This allowed the light-sheet to illuminate a 1.46mm wide region in sample space. In the other dimension, the beam was truncated by an adjustable slit, and then focused by a cylindrical lens into a light-sheet. The light-sheet was focused on a resonant galvo mirror for shadow suppression, which was conjugate to the focal plane of the illumination objective. By adjusting the size of the slit, the effectively used numerical aperture of the excitation objective could be varied.

For the high-resolution mode using axially swept light-sheet microscopy (ASLM), as shown in **Supplementary Figure 2**, the beam was first expanded 20-fold. The beam was then shaped by a cylindrical lens into a sheet, which was focused on resonant galvo. This galvo was conjugate to the focal plane of the remote focusing objective. In the focal plane of the remote focusing objective, the laser light was back-reflected by a mirror. The back-reflected light was then mapped into the illumination objective, whose full numerical aperture was leveraged to focus the light-sheet into the sample plane. The focal length of the illumination objective varies with immersion media, the value indicated applies for water. The half angle $\alpha$ of the objective is $16^o$, and as such, the NA also varies with immersion index (NA=n*sin($\alpha$) ). For water, the NA is 0.367.

We used the following equations[1] to estimate the thickness and propagation length of the light-sheet in sample space:

The thickness can be estimated by the well-known Abbe equation:

$$d_{thickness} = \lambda / 2NA \ .$$

The effective length of the light-sheet can be estimated by:

$$d_{prop} = \frac{\lambda}{n * (1 - \cos\alpha)} \ .$$

$\lambda$ is the excitation wavelength, NA is the numerical aperture and $\alpha$ is the half opening angle of the objective, here $16^o$. For the high-resolution mode, we found a light-sheet thickness of ~664nm for 488nm excitation. The propagation length was estimated to 9.5 microns. In the low-resolution mode, the light-sheet could be continuously adjusted by opening and closing the motorized slit aperture.

To make the ALSM mode work, the camera required a synchronized rolling pixel readout, which depending on the manufacturer is implemented and notated differently. In **Supplementary Figure 3** we show the implementation on the Photometrics camera.

**Supplementary Table 1: Hardware components of the multi-scale microscope**. The table is divided into components for illumination (laser, LED: orange), illumination path (white, shared between both illumination paths), components of the low-resolution illumination path (green) and high-resolution illumination path (blue), sample chamber (violet), detection path (gray) and diverse and computer (white).

| Category | Item | Supplier / Catalog Number |
|---|---|---|
| Laser | OBIS 488nm LS 100mW LASER | Coherent, 1226419 |
| Laser | OBIS 552nm LS 100mW LASER | Coherent, 1230941 |
| Laser | OBIS 594nm LS 100mW LASER | Coherent, 1285743 |
| Laser | OBIS 640nm LX 100mW LASER | Coherent, 1185055 |
| Laser | Laser Box: OBIS: 2K Ohm: 5 Bay with Power Supply and USA Power Cord | Coherent, 1343229 |
| LED illumination | Arduino UNO REV3 | Arduino, A000066 |
| LED illumination | Clear White LED | Microtivity, IL188, SYNCHKG022865 |
| Illumination path | Mirrors | Thorlabs, PF10-03-P01- Ø1" Protected Silver Mirror |
| Illumination path | Dichroic, 613 nm edge LaserMUX™ single-edge laser dichroic beamsplitter | Semrock, LM01-613-25 |
| Illumination path | Dichroic, 573 nm edge BrightLine® single-edge standard epi-fluorescence dichroic beamsplitter | Semrock, FF573-Di01-25x36 |
| Illumination path | Dichroic, 503 nm edge LaserMUX™ single-edge laser dichroic beamsplitter | Semrock, LM01-503-25 |
| Illumination path | f=50 mm, Ø1" Achromatic Doublet | Thorlabs, AC254-050-A-ML - f=50 mm |
| Illumination path | Pinhole, 40 um | Thorlabs, P40D |
| Illumination path | f=200 mm, Ø1" Achromatic Doublet | Thorlabs, AC254-200-A-ML, - f=200 mm |
| Illumination path | Motorized rotation stage for half waveplate | Thorlabs, ELL14K |
| Illumination path | Halfwave plate | Bolder Vision Optik, Inc, BVO AHWP3 (λ/2) |
| Illumination path | Cube Beamsplitter, Polarizing, 25.4 mm, 420-680 nm | NewPort,10FC16PB.3 |
| Illumination path – low resolution | Motorized flip mirror to block low resolution illumination during high-res imaging | Thorlabs, MFF101 - Motorized Filter Flip Mount with Ø1" Optic Holder, 8-32 Tap |
| Illumination path – low resolution | 25 mm Achromatic cylindrical lens | Edmund Optics, Stock #68-160 |
| Illumination path – low resolution | 100 mm Cylindrical Achromatic Doublets | Thorlabs, ACY254-100-A - f = 100.0 mm |
| Illumination path – low resolution | Adjustable Mechanical Slit | Thorlabs, VA100 |

| | | |
|---|---|---|
| Illumination path – low resolution | 200 mm Cylindrical Achromatic Doublets | Thorlabs, ACY254-200-A - f = 200.0 mm |
| Illumination path – low resolution | Resonant galvanometer for m-SPIM illumination | Cambridge Technology, CRS 4 KHz with UCTRONICS DC 0-10V 0/4-20mA Current Voltage Signal Generator and Acopian Power Supply Model A12MT400 |
| Illumination path – low resolution | f=100 mm Ø2" Achromatic Doublet | Thorlabs, AC508-100-A-ML - f=100 mm |
| Illumination path – low resolution | Illumination objective, nominal NA of 0.4, WD of 12 mm | Applied Scientific Instrumentation, 54-10-12 Multi-Immersion Objective |
| Illumination path – high resolution | Halfwave plate | Bolder Vision Optik, Inc, BVO AHWP3 (λ/2) |
| Illumination path – high resolution | 5x Beam Expander | Thorlabs, 5X Achromatic Galilean Beam Expander, AR Coated: 400 - 650 nm |
| Illumination path – high resolution | Motorized mechanical slit | Standa, 10MAOS10-1 with XIMC Motor Controller |
| Illumination path – high resolution | 50 mm Cylindrical Achromatic Doublets | Thorlabs, ACY254-50-A - f = 50.0 mm |
| Illumination path – high resolution | Resonant galvanometer for m-SPIM illumination | Cambridge Technology, CRS 4 KHz with UCTRONICS DC 0-10V 0/4-20mA Current Voltage Signal Generator and Acopian Power Supply Model A12MT400 |
| Illumination path – high resolution | f=200 mm Ø2" Achromatic Doublet | Thorlabs, AC508-200-A-ML - f=200 mm |
| Illumination path – high resolution | Cube Beamsplitter, Polarizing, 25.4 mm, 420-680 nm | NewPort,10FC16PB.3 |
| Illumination path – high resolution | Quarterwave plate | Bolder Vision Optik, Inc, BVO AQWP3 (λ/4) |
| Illumination path – high resolution | Remote focus objective | Olympus, XLFLUOR4X/340 |
| Illumination path – high resolution | 25mm Dia., 3mm Thick, VIS 0° Coated λ/4 N-BK7 Window | Edmund Optics, BK7 Glass 37-005 |
| Illumination path – high resolution | Linear focus actuator | Equipment Solutions, LFA2010 stage set with SCA814 and SPS15 servo-controlled amplifier |
| Illumination path – high resolution | f=200 mm Ø2" Achromatic Doublet | Thorlabs, AC508-200-A-ML - f=200 mm |
| Illumination path – high resolution | f=75 mm Ø2" Achromatic Doublet | Thorlabs, AC508-75-A-ML - f=200 mm |
| Illumination path – high resolution | Illumination objective, nominal NA of 0.4, WD of 12 mm | Applied Scientific Instrumentation, 54-10-12 Multi-Immersion Objective |
| Sample Chamber | Custom design | Manufactured with Protolabs |
| Sample Chamber | DC power supply for heating | MPJA, 9312-PS |
| Sample Chamber | Digital Temperature Switch with temperature switch probes | Dwyer, TS2-011<br>Dwyer, TS-7 |
| Sample Chamber | Silicone Rubber Fiberglass Flexible Heater 450°F Max to 48" | Omega, SRFGA-203/10-P |

| | | |
|---|---|---|
| Sample Chamber | Rotation Stage | SmarAct, SR-2812 with MCS controller |
| Sample Chamber | Linear Piezo Stage with 83 mm travel range | SmarAct, SLC-24120 - Linear Piezo Stage |
| Sample Chamber | Linear Piezo Stage with 21 mm travel range | SmarAct, SLS-3232 with MCS2 controller, with Ethernet port, for all three stages |
| Detection Path | Detection objective, 20x Objective with NA1.0, 2 mm WD | N20X-PFH - 20X Olympus XLUMPLFLN Objective |
| Detection Path | Motorized Flip Mirror | Newport, 8893-K |
| Detection Path | Thorlabs, Piezo Stage for detection objective alignment | PIA25, 25 mm Travel Range, with KIM001 controller |
| Detection Path | Filter wheel, 6 positions, 25mm | Ludl Electronics, 96A351 with MAC6000 controller, 996066. |
| Detection Path | Emission Filter for 488 nm excitation | Semrock, FF01-525/30-25 |
| Detection Path | Emission Filter for 552 nm excitation | Semrock, FF01-572/28-25 |
| Detection Path | Emission Filter for 594 nm excitation | Semrock, FF02-615/20-25 |
| Detection Path | Emission Filter for 640 nm excitation | Semrock, FF01-676/37-25 |
| Detection Path | Tube lens 100 mm, low-resolution detection path | AC508-100-A-ML - f=100 mm, Ø2" Achromatic Doublet |
| Detection Path | Tube lens 500 mm, high-resolution detection path | ACT508-500-A-ML - f=500 mm, Ø2" Achromatic Doublet |
| Detection Path | Low-resolution camera | Teledyne Photometrics, Iris 15, 4.25 µm x 4.25 µm pixel size |
| Detection Path | High-resolution camera | Teledyne Photometrics, Prime BSI Express, 6.5 µm x 6.5 µm pixel size |
| Microscope computer | Analog Output Device | NI PCIe-6738 |
| Microscope computer | Microscope control computer | Supermicro, SYS-7049GP-TRT, Windows 10 Enterprise, Intel Xeon Silver 4112 CPU @2.6GHz, 96 GB RAM, data drive: INTEL SSDPE2KE076T8 and operating system drive: INTEL SSDPE2KX010T8, NVIDIA Quadro P4000 graphics card, Intel Ethernet Controller X550 network card, |
| Diverse | Diverse BNC cables | |
| Diverse | Diverse optomechanical mounts, irises and | |
| Diverse | Diverse USB-HUB, Ethernet port, RS-232 hub | |

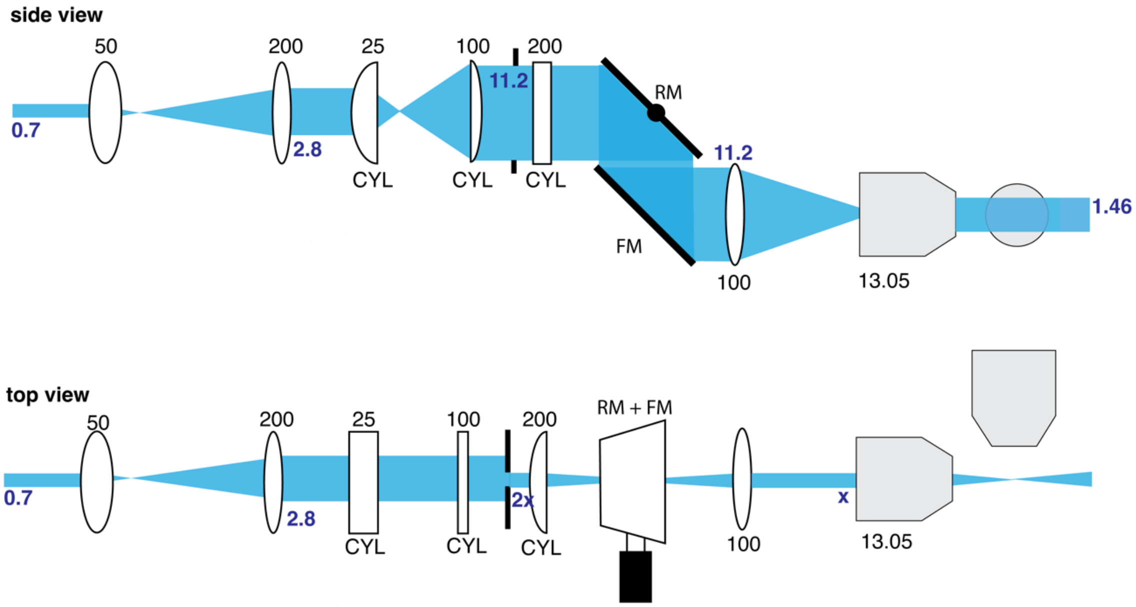


**Supplementary Figure 1: Schematic of the left low-resolution arm of the dual-sided multi-scale microscope illumination.** Numbers in black indicate the focal length of the corresponding lens. Numbers in blue approximate the beam size. CYL cylindrical lens, FM folding mirror, RM resonant galvo mirror

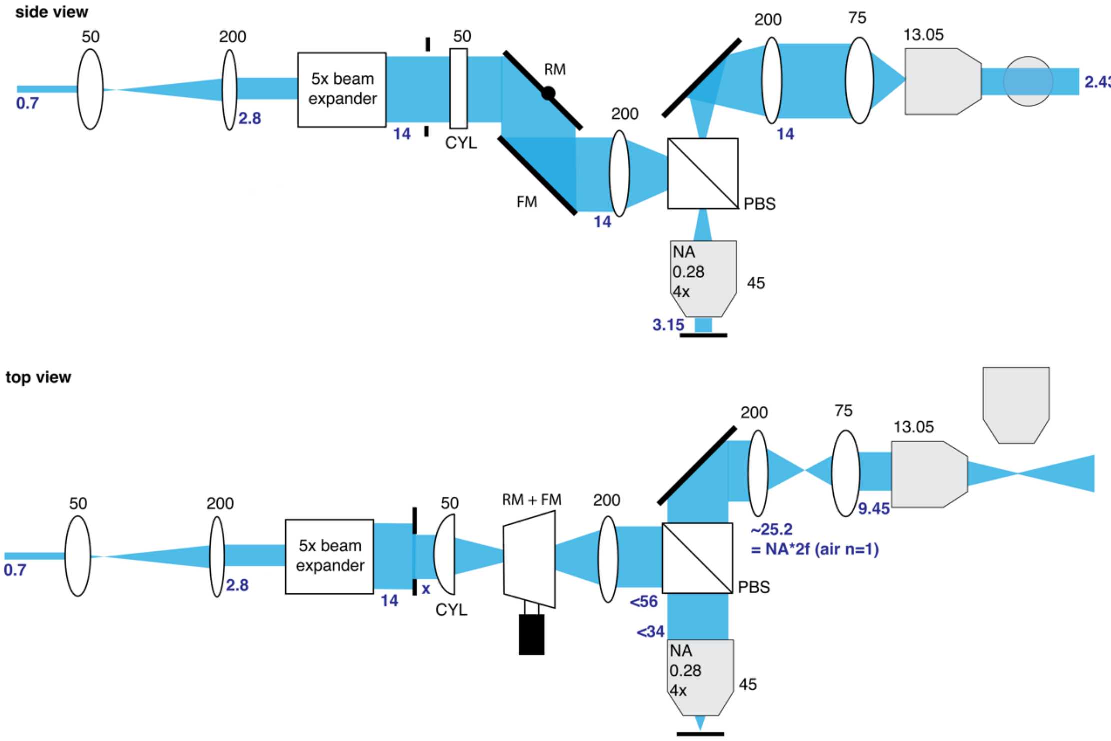


**Supplementary Figure 2: Schematic of the high-resolution arm of the dual-sided multi-scale microscope illumination**. Numbers in black indicate the focal length of the corresponding lens. Numbers in blue approximate the beam size. CYL cylindrical lens, FM folding mirror, RM resonant galvo mirror, PBS polarizing beam splitter.

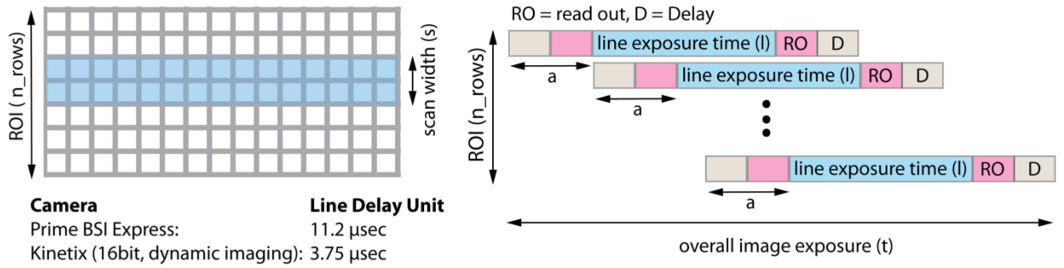


**Formula derivation:**

(i) s = l / a => a = l/s; with s = scan width, a = sum of delay and read out, l = line exposure time
(ii) t = ROI * a + l + a = (ROI +1) * a + l; with ROI = number of rows
Insert (i) into (ii): t = (ROI+1) *l/s + l and solve for l: (iii) l = t / ((ROI +1)/s +1)
Now, calculate the delay factor parameter which is needed for the programmable scan mode.
(iv) a = read out + delay = (line delay unit) * (delay factor +1)
Use (ii) to obtain: a = (t-l)/(ROI+1) and insert (iv): (v) delay factor = (t-l)/[(ROI+1) * line delay unit] - 1

(iii) $$\text{line exposure time} = \frac{\text{overall image exposure}}{1 + (\text{ROI}+1) / (\text{scan width})}$$

(v) $$\text{delay factor} = \text{ceil}\left(\frac{\text{image exposure time - line exposure time}}{(\text{ROI} +1) * \text{line delay unit}}\right) - 1$$

Now recalculate the exposure time as you need to provide integers (ceil) the functions

$$\text{overall image exposure} = \text{line exposure time} + (\text{ROI} +1) * (\text{line delay unit}) * (\text{delay factor} +1)$$

**Supplementary Figure 3: Axially Swept Light-Sheet imaging with Photometrics**. Formulas to calculate the parameters for Axially Swept Light Sheet (ASLM) imaging with the programmable scan mode on a Photometrics camera.

**Supplementary Note 2: Fluorescent Nanospheres measurements**

We estimated the resolution of the microscope modalities by acquiring 3D stacks of fluorescent nanospheres embedded in Agarose, and measuring the Full width half maximum in the x, y and z direction on multiple nanosphere images. To determine whether the measured resolution values (n=20, each condition) followed a normal distribution, we visualized the measurements with Q-Q plots (**Supplementary Figure 5**) and applied the Shapiro-Wilk test. The results (p-values) of the Shapiro-Wilk test were as follows:

- Low-resolution mSPIM PSF: lateral (x): 0.5691, (y): 0.1107; axial (z): 0.3771
- High-resolution mSPIM PSF: lateral (x): 0.0524, (y): 0.2555; axial (z): 0.9628
- High-resolution ASLM PSF: lateral (x): 0.5677, (y): 0.2021; axial (z): 0.0581

Thus, the data provided no evidence that the measured beads are significantly different from a normal distribution.

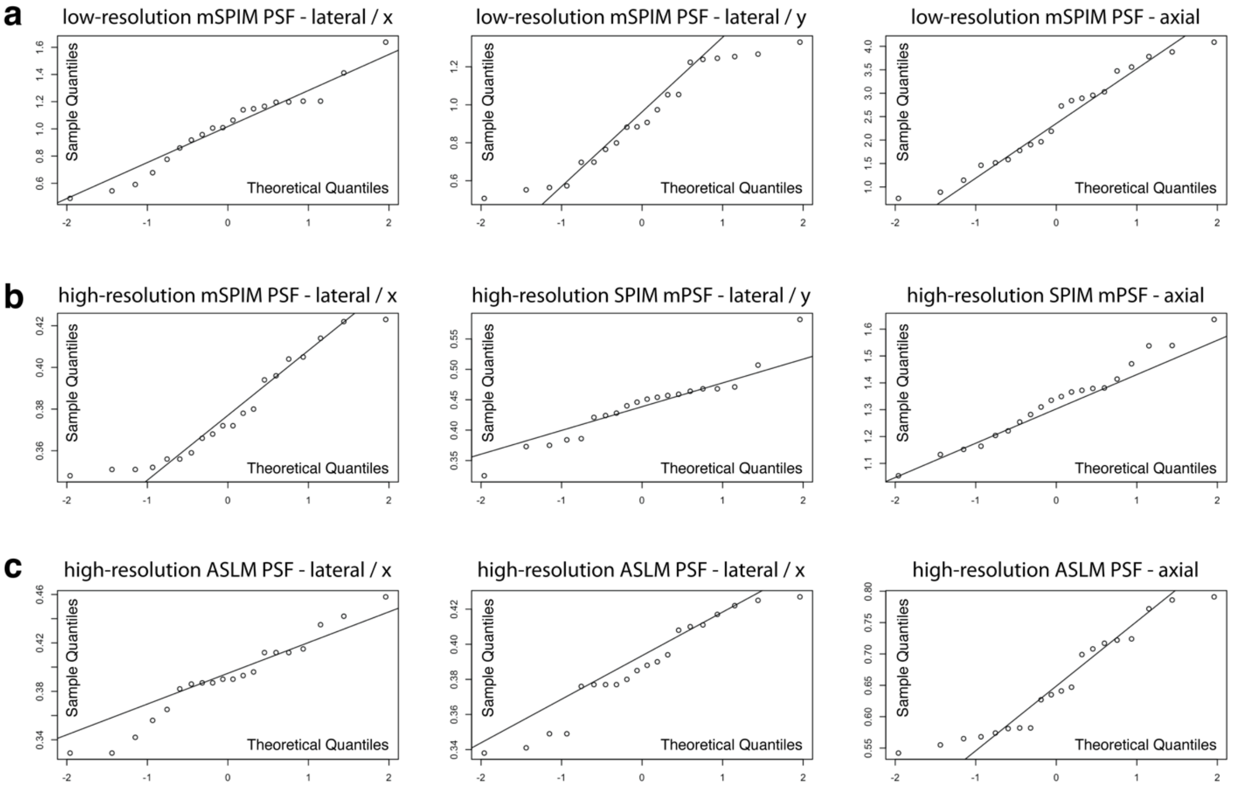


**Supplementary Figure 5: Bead measurement data**. Quantile-quantile plot of bead measurements in the different microscope resolution levels: analysis of **a** low-resolution SPIM mode PSF values, **b** high-resolution SPIM mode PSF values, and **c** high-resolution ASLM PSF mode PSF values. N=20 for each panel.

In **Supplementary Figure 4,** images of beads in Agarose using the different image modalities, as well as Full width half maximum measurements are shown. The top row shows the point spread function in the low-resolution mode, that is low-resolution light-sheet and low-resolution detection path. By switching the detection to high resolution and opening the motorized slit, a laterally tight PSF emerges, which is however still axially extended to over one micron width. When switching to the ASLM mode, the axial resolution is improved to about 650nm, which is in correspondence to our estimate of the light-sheet's thickness in that modality (**Supplementary Note 1**).

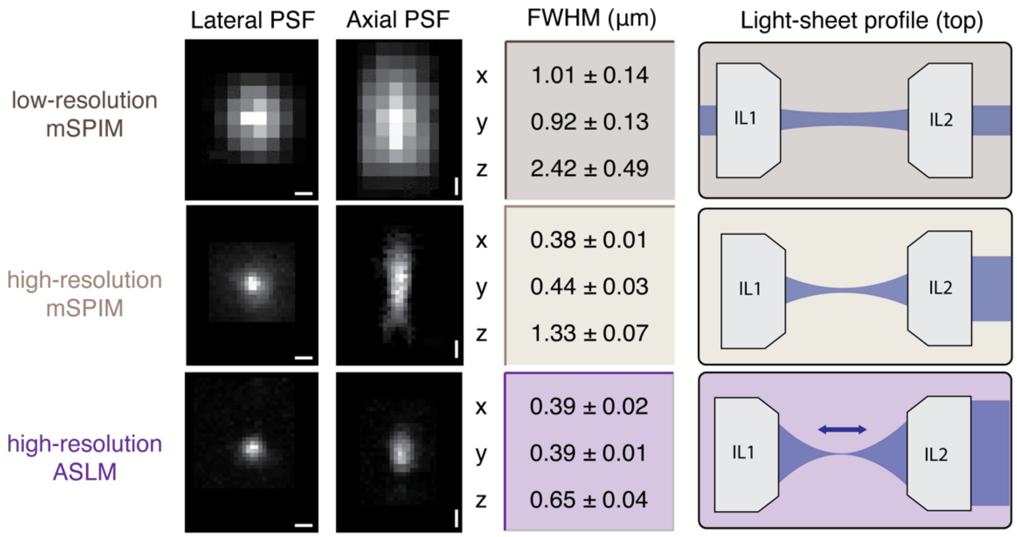


**Supplementary Figure 4: Comparison of all acquisition modes.** Comparison of all available modalities on the multi-scale microscope, including low-resolution multi-directional selective plane illumination microscopy (mSPIM), high-resolution mSPIM and high-resolution axially swept light-sheet microscopy (ASLM). Representative PSF images of 0.2 $\mu$m YG Microspheres are displayed with calculated FWHM values of the beads (n=20 beads each). The last column schematically depicts the light-sheet profile (top view) to highlight the difference between the modalities. The dark blue arrow indicates the scan direction of the thin light-sheet. Scale-bar lengths are as follows: 0.5 µm.

#### Supplementary Note 3: Microscope Control Software Architecture

In this note, we provide more detail about the microscope hardware control software. In **Supplementary Figure 6**, we provide waveforms required for a 3D stack acquisition using axially swept light-sheet microscopy (ASLM). These waveforms are generated in the Python file *acquisition_array_class.py* as NumPy arrays and sent to the NI Data acquisition card for precise timing of all components. In **Supplementary Figure 7**, we provide an overview about the organization of the microscope acquisition code with its separation into an MVC design pattern: Model (*multiScope.py,* and the corresponding subfolders), View (gui folder), and Controller (*multiScale_main.py*). **Supplementary Figure 8** highlights the intuitive graphical user interface for controlling and running long-term acquisitions on the microscope.

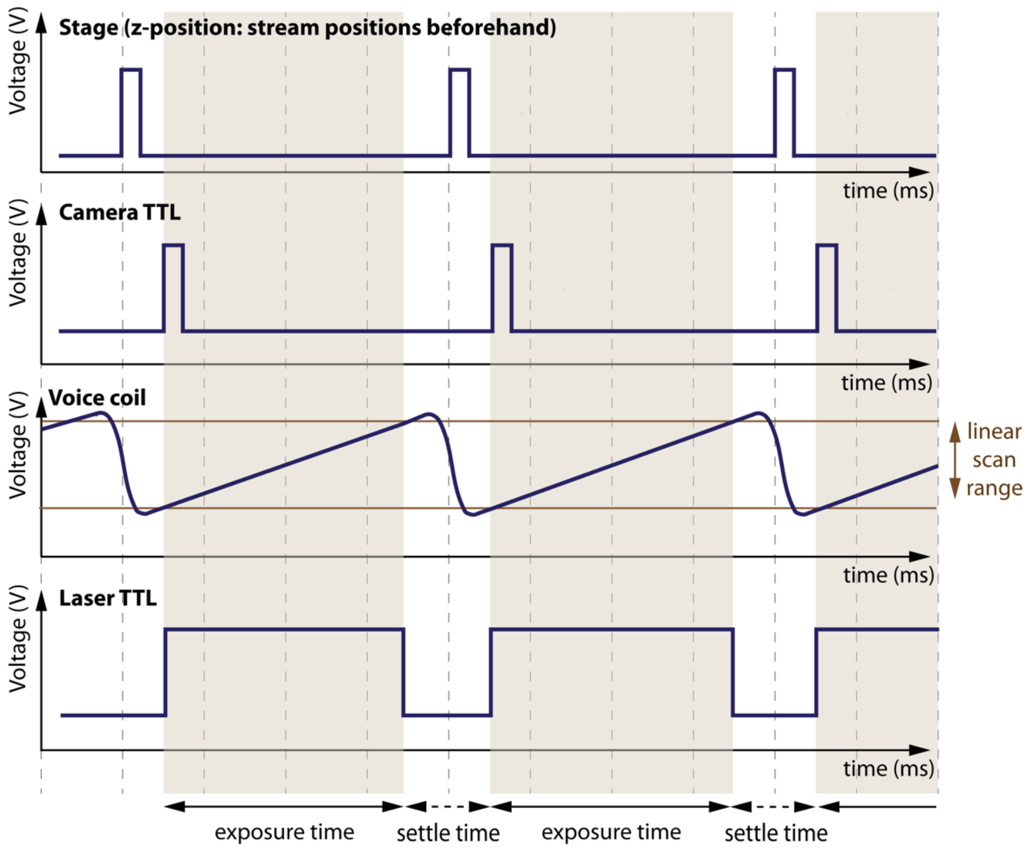


**Supplementary Figure 6: Acquisition waveforms for hardware timing control.** To control the precise timing of different hardware components on the microscope, an NI DAQ card was used to send trigger signals and voltage information to the devices. Thereby, for an Axially Swept Light-Sheet (ASLM) volumetric acquisition, the stage, camera, voice coil and laser needed to be synchronized.

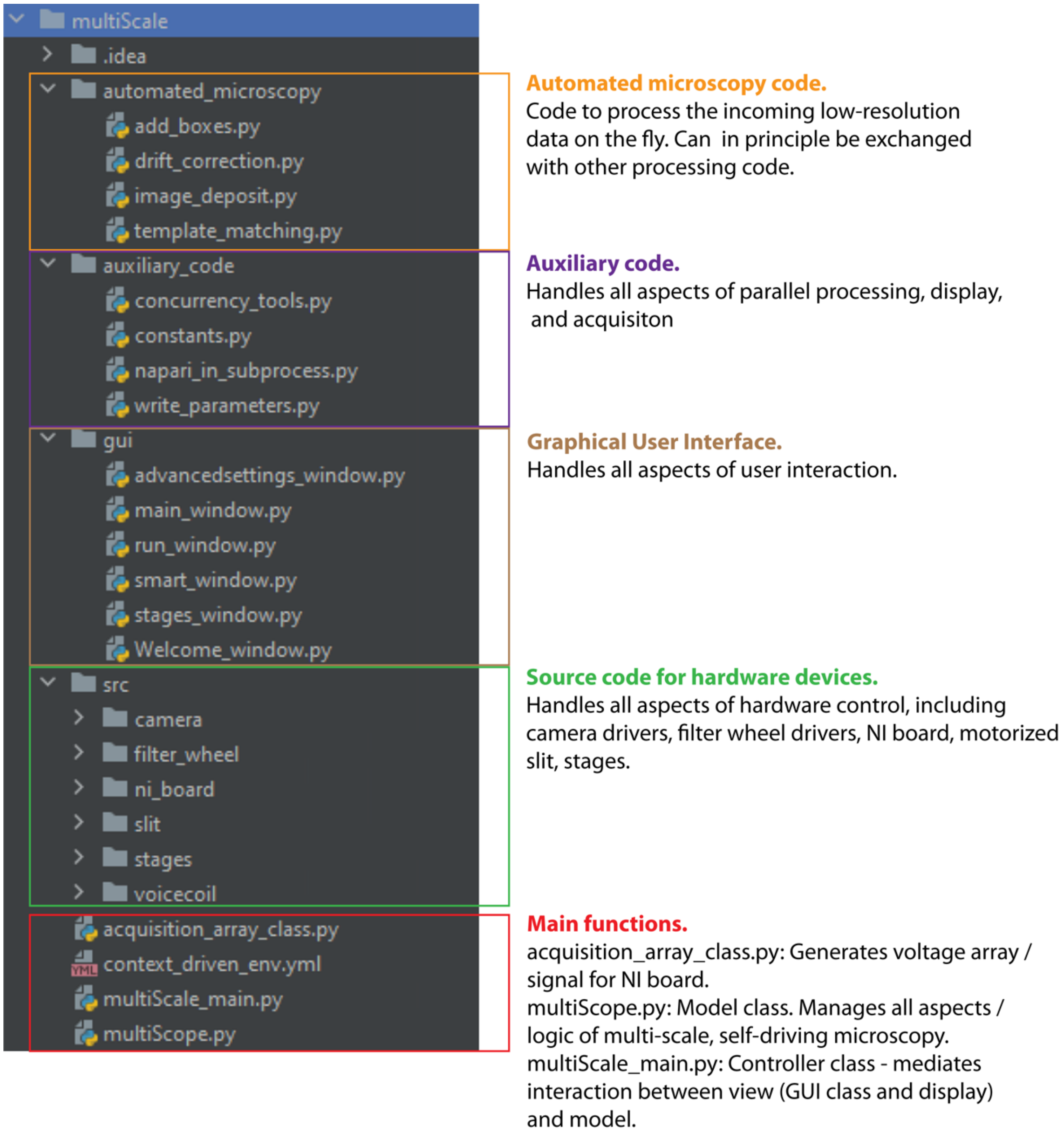


**Supplementary Figure 7: Self-driving, multi-scale control software.** Overview over the MVC design and file structure of the microscope control software.

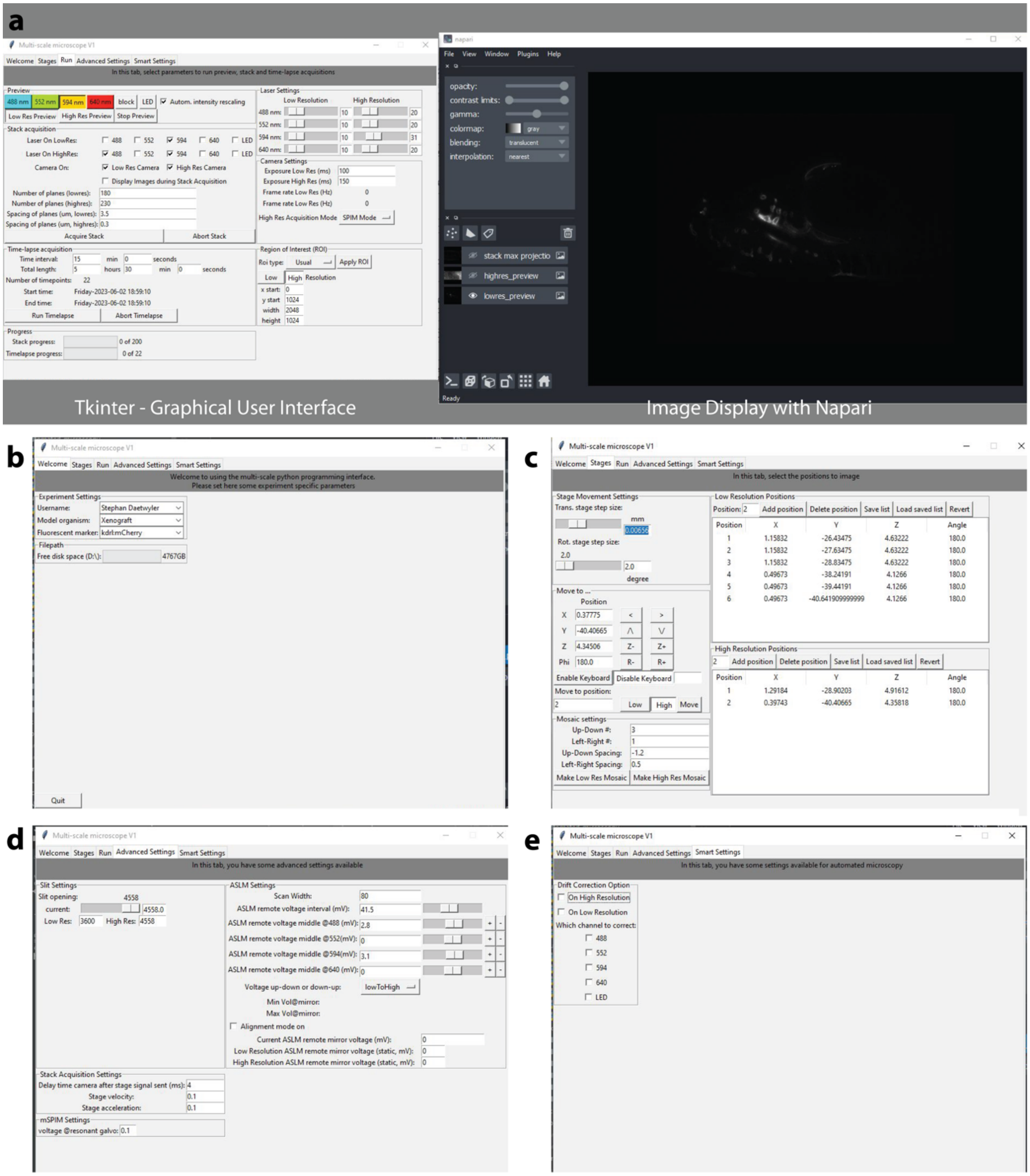


**Supplementary Figure 8: Overview over the graphical user interface and display of the microscope control software**. **a** Image display with Napari and "Run tab" to start a preview, stack and time-lapse acquisition with camera, laser and region of interest settings. **b** Start up window to control folder name settings, **c** Stage settings to select imaging positions and move the stages, **d** Advanced settings to control ASLM parameters, motorized slit, and time delays for stack acquisition, **e** Self-driving microscope settings.

**Supplementary Note 4: A375 xenograft experiments**

In this note, we provide more detail about the xenograft experiments with A375 melanoma cancer cells. In **Supplementary Figure 9**, we show a representative time-lapse experiment of how macrophages are in close contact with A375 cells, but not inducing phagocytosis. Instead, we observed a cell division over the course of observation. This is also reflected in the high survival rate of cancer cells in zebrafish tail metastasis (**Supplementary Figure 10**). We have however observed apoptosis events of A375 cells in the tail without initial involvement or close attachment of macrophages (**Supplementary Figure 11**). However, macrophages cleared the apoptotic bodies after cancer cell apoptosis (**Supplementary Figure 11, 12**).

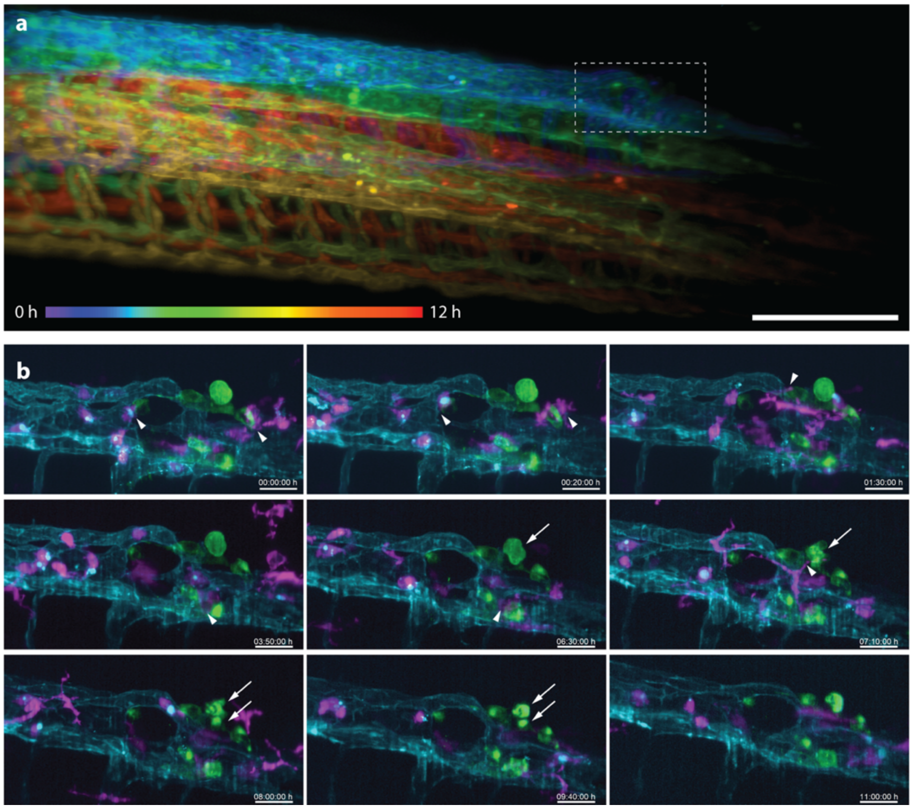


**Supplementary Figure 9: A375 cells interacted with macrophages but were not phagocytosed.** **a** Low-resolution imaging of the vasculature, labeled with Tg(*kdrl:Hsa.HRAS-mCherry*)[2], revealed the large displacement of the zebrafish tail over the course of 12 hours of imaging, highlighting the need for self-driving multi-scale microscopy of a region of interest (dashed box). The sample was imaged every 10 min and selected maximum intensity projections (every 1:30 hours) were color coded and overlaid onto each other. **b** To reveal cell-cell interactions, we employed high-resolution ASLM imaging (Supplementary Movie 7) of a selected region of interest with a cluster of A375 melanoma cancer cells (green, pVimentin-PsmOrange label). Selected maximum intensity projections of the time-lapse experiment are displayed. For tissue context, the vasculature (cyan) was labeled with Tg(*kdrl:Hsa.HRAS-mCherry*)[2]. Over the time-lapse experiment, we observed cell division of the A375 cells (white arrow). Interestingly, macrophages (magenta), labeled with *Tg(mpeg1:EGFP)*[3], interacted with cancer cells (white arrowheads). However, despite these multiple interactions, none of the A375 cells was phagocytosed, hinting at mechanisms of immune evasion. Scale-bar lengths are as follows: **a** 200 µm; **b** 30 µm.

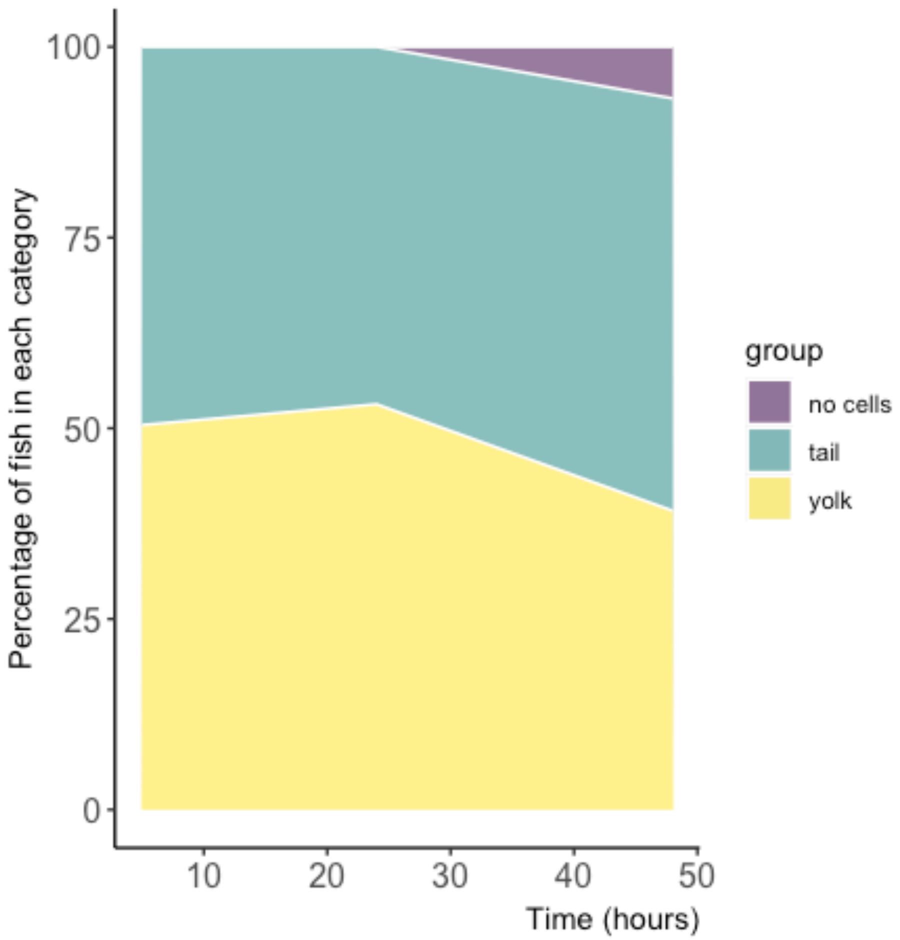


**Supplementary Figure 10: Survival curve of A375 melanoma cells.** We quantified survival of A375 melanoma cells in a larval xenograft assay over 48 hours post injection (hpi) (n=117 fish at 0 hpi). The different categories indicate presence of metastasis in the tail (cyan) or only at the injection site (yolk, yellow). Zebrafish without cells are labeled as "no cells" (violet). To compare time points, the percentage of all surviving fish are displayed in each category.

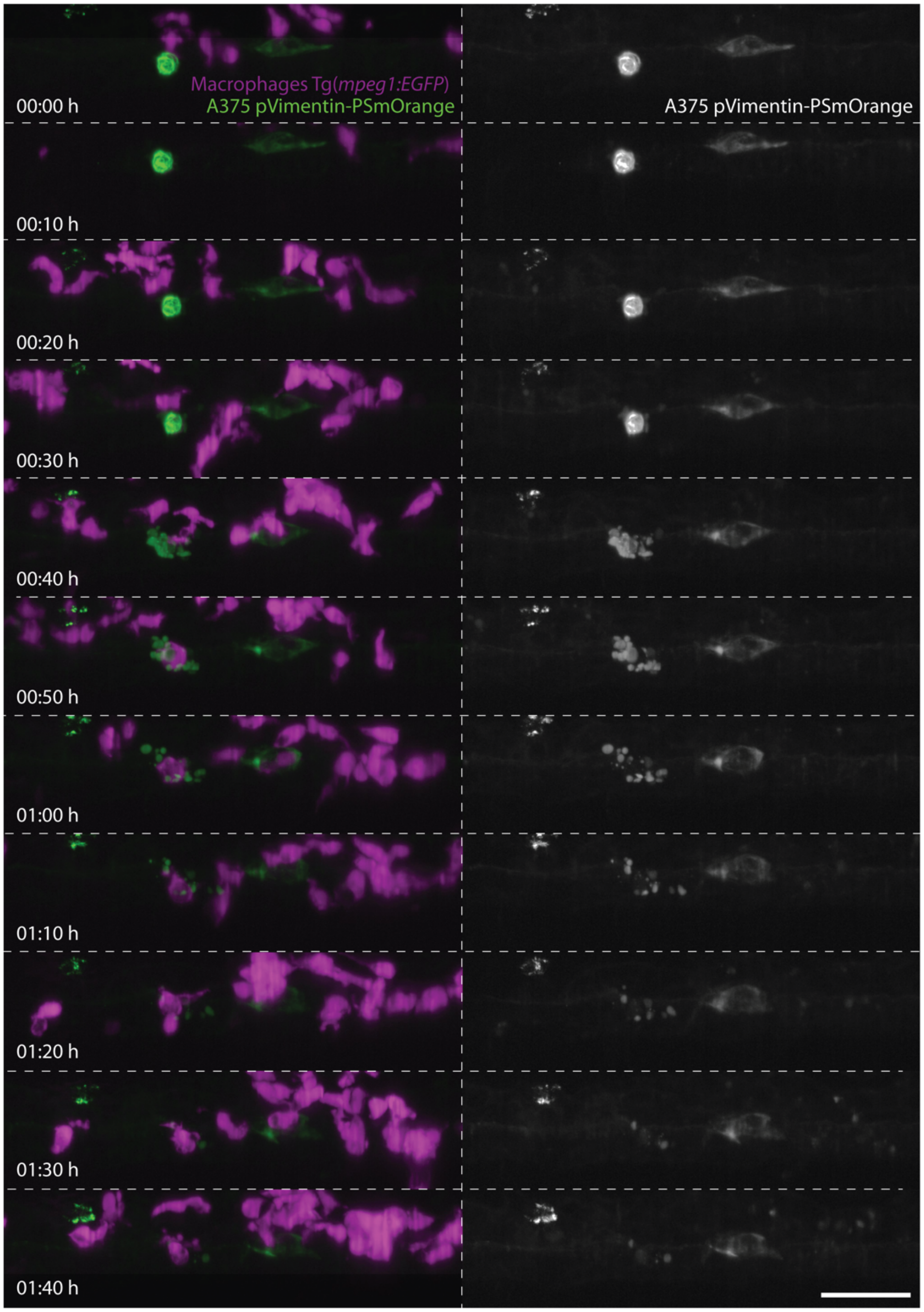


**Supplementary Figure 11: Apoptosis of A375 cells and subsequent clearance of cell debris by a macrophage.** A time-lapse experiment using self-driving high-resolution microscopy revealed how a A375 melanoma cancer cell, labeled with pVimentin-PsmOrange (left panel in green, right in gray), underwent apoptosis and formed apoptotic bodies that were then cleared by a zebrafish macrophage, labeled with *Tg(mpeg1:EGFP)*[3] (left panel in magenta). Scale-bar lengths are as follows: 50 µm

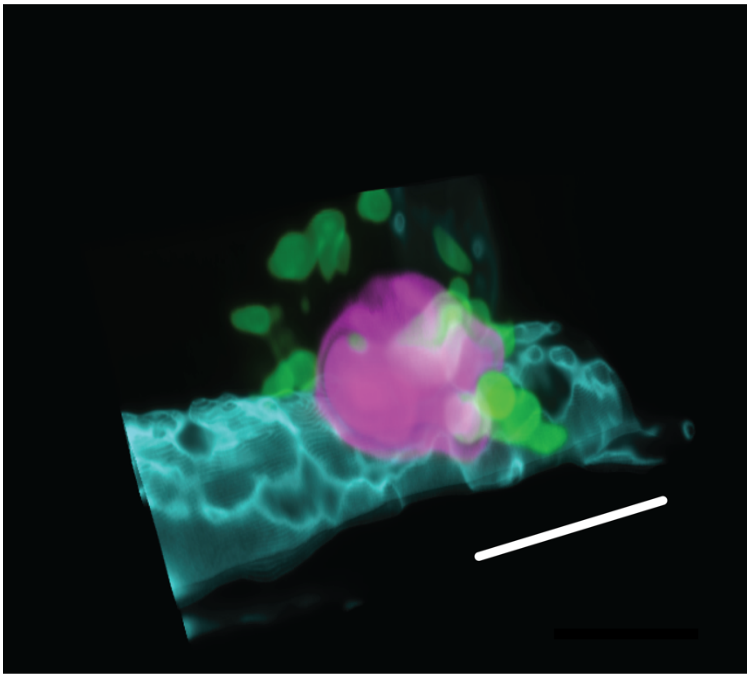

**Supplementary Figure 12: 3D rendering of a macrophage taking up apoptotic bodies of an A375 cells.** The rendering displays a single timepoint at 00:50 h from Supplementary Figure 11 above. Apoptotic bodies of an A375 cells (green), labeled with pVimentin-PsmOrange, remained after the cell underwent apoptosis. They were cleared by a macrophage (magenta), labeled with *Tg(mpeg1:EGFP)*[3]. For tissue context, the vasculature (cyan), labeled with Tg(*kdrl:Hsa.HRAS-mCherry*)[2] is displayed. Data was rendered with AGAVE[4]. Scale-bar lengths are as follows: 20 µm

**Supplementary Note 5: Data analysis**

In this note, we provide more details about the analysis of multi-scale data. This includes a detailed description of the low-resolution macrophage segmentation pipeline, based on py-clesperanto and CLIJ[5] (**Supplementary Figure 13**). Moreover, we describe how we calculated the zebrafish volume to simulate random cell distribution within the organisms for the Hopkins statistics[6,7] calculation (**Supplementary Figure 14**), and provide a list of all global morphological features used for the calculation of the shape morphological space of the high-resolution data (**Supplementary Table 2**). Additionally, we provide a more detailed analysis of the PCA space that was established to compare macrophage morphologies over time (**Supplementary Figure 15**).

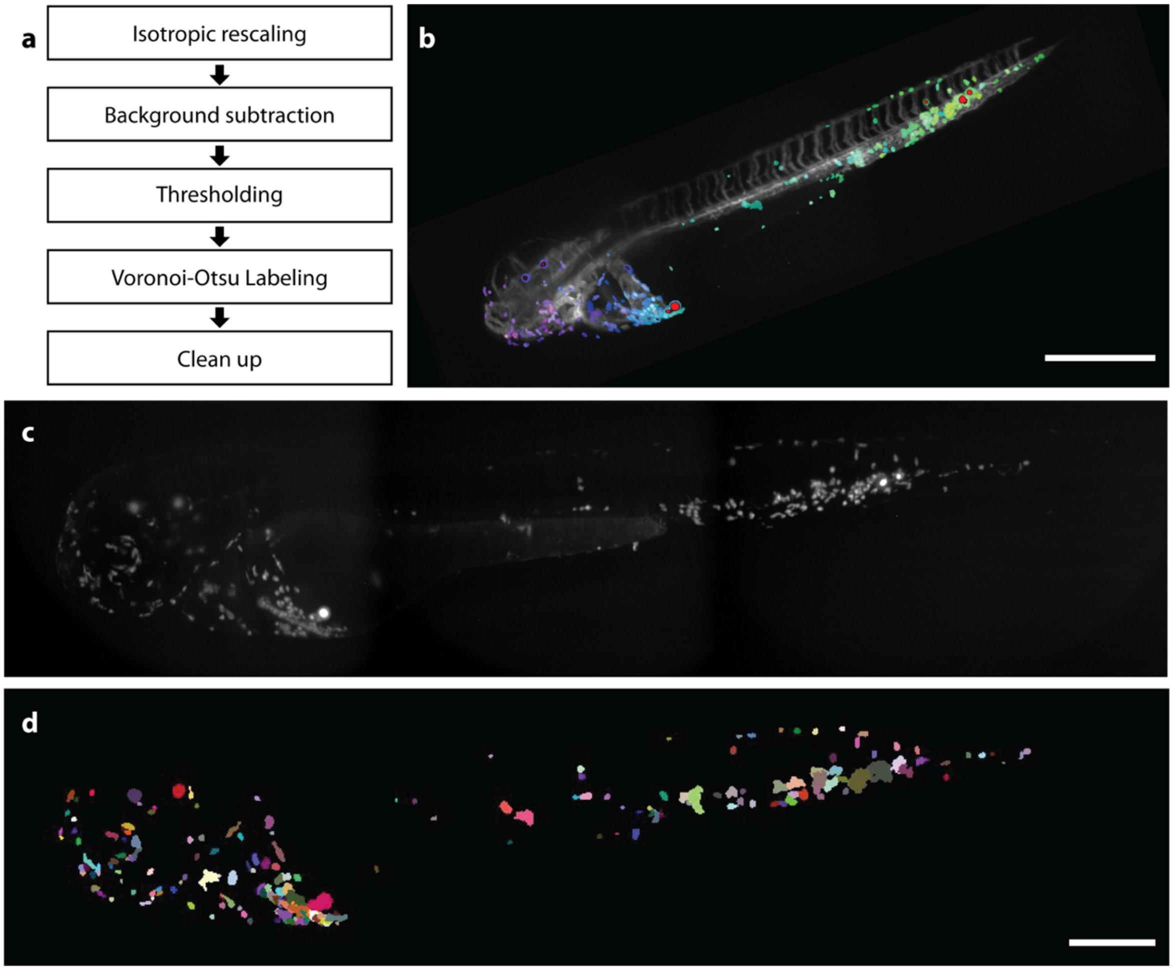


**Supplementary Figure 13: Workflow to obtain a low-resolution segmentation of zebrafish macrophages**. **a** Schematic description of segmentation pipeline using the py-clesperanto library. **b** 3D rendering of resulting segmentation (color coded from head to tail in violet to green) of macrophages, labeled with *Tg(mpeg1:EGFP)*[3] and U-2 OS cancer cells (red), labeled with pVimentin-PSmOrange. For tissue context, the vasculature, labeled with Tg(*kdrl:Hsa.HRAS-mCherry*)[2], was displayed in gray. **c** Maximum intensity projection of the raw data (488 nm excitation) of macrophages (gray) and cancer cell bleed-through (bright white) across whole zebrafish larva. **d** The corresponding maximum intensity projection of the macrophage segmentation (in color). Scale-bar lengths are as follows: **b** 500 µm; **c,d** 250 µm.

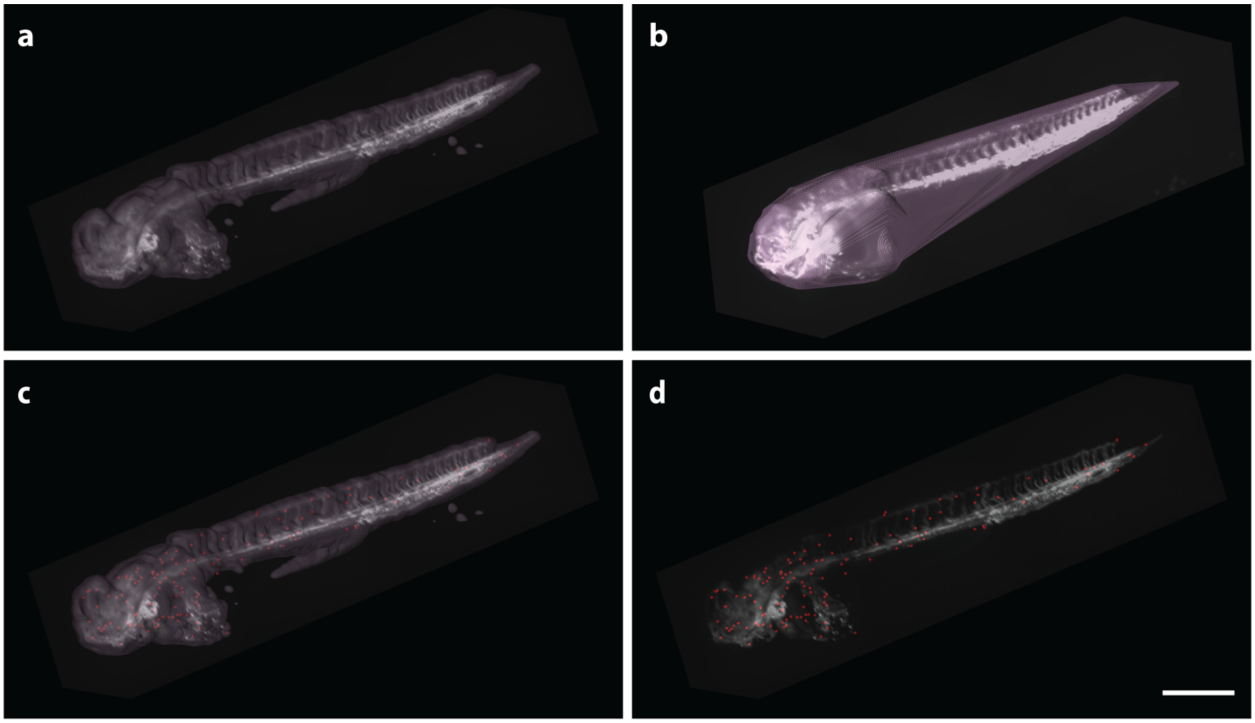


**Supplementary Figure 14: Simulation of random macrophage distribution inside zebrafish.** To simulate points inside the zebrafish for analysis of cell dissemination, we calculated the fish outline based on the stitched low-resolution vascular channel Tg(*kdrl:Hsa.HRAS-mCherry*)[2]. **a** 3D rendering of the fish outline (violet) overlaid onto the vascular channel (gray). To obtain it, we first Gamma corrected (gamma=0.9) and smoothed the vascular data with a 3D Gaussian kernel (σ = 10,10,3 pixels). Then, as we estimated the fish volume to be roughly 10% of the 3D stack, we calculated the threshold value by analyzing the stack histogram, and selected the pixels with the highest 10% intensity values. Next, we performed a hole filling morphological operation on the segmented fish and expanded the segmentation by smoothing with a 3D Gaussian kernel (σ = 20,20,5 pixels) followed by another thresholding (threshold at 5). After another hole filling operation, we obtained the fish outline. To calculate the fish outline, we used a macro script in ImageJ/Fiji[8] . The script is available on GitHub (*generate_fish_outline_forSegmentation_percentile.ijm)*. **b** Based on the outline in a, we could also calculate the convex hull (violet). However, after comparing the analysis using the complex hull against the outline based on the segmentation, we did not see any difference. **c** Therefore, we used the outline (violet) to simulate macrophage dissemination (red) randomly in the zebrafish larvae. The number of simulated macrophages corresponded to the number of segmented macrophages per time point. **d** 3D rendering of simulated macrophages (red) overlayed on the vascular channel. Scale-bar length: 500 µm

**Supplementary Table 2:** Global geometric features used to quantify and compare macrophage morphology

| **Global geometrical features** | **Definition** |
|---|---|
| Volume (V) | |
| Surface area (S) | |
| Solidity | $V_{\mathrm{Cell}} / V_{\mathrm{Convex\ hull}}$ |
| Sphericity | $\pi^{1/3} (6V_{\mathrm{Cell}})^{2/3} / S_{\mathrm{Cell}}$ |
| Long length | Longest length in all direction |
| Extend | $V_{\mathrm{Cell}} / V_{\mathrm{Bounding\ box}}$ |
| Aspect ratio | $L_{\mathrm{Short}} / L_{\mathrm{Long}}$ |
| Roughness | $S_{\mathrm{Cell}} / S_{\mathrm{Convex\ hull}}$ |
| Volume sphericity | $V_{\mathrm{Cell}} / V_{\mathrm{Cicumscribed\ sphere}}$ |
| Radius sphericity | $\mathrm{Equivalent\ radius} / R_{\mathrm{Cicumscribed\ sphere}}$ |
| Ratio sphericity | $R_{\mathrm{Inscribed\ sphere}} / R_{\mathrm{Cicumscribed\ sphere}}$ |
| Circumscribed sphere area ratio | $S_{\mathrm{Cell}} / S_{\mathrm{Cicumscribed\ sphere}}$ |

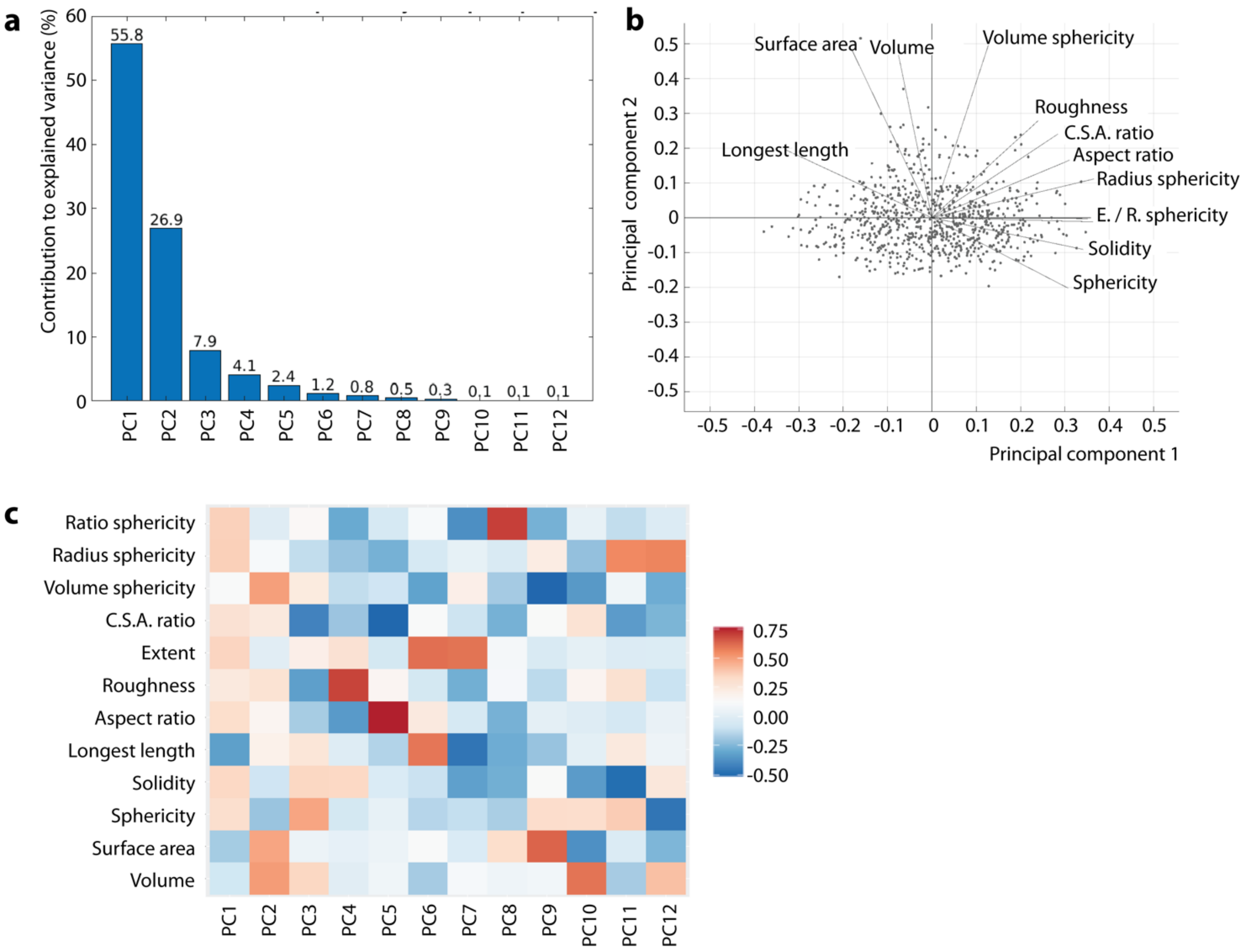


**Supplementary Figure 15: PCA analysis of macrophage shapes. a** The first two principal components explained over 80% of the observed variance in the dataset. **b** To understand, which features contributed to the first two principal components, we generated the biplot of the principal component analysis, and **c** calculated the Z-normalized correlation matrix between the dataset and PC-space. Abbreviations: PC Principal Component; E. Extend; C.S.A. Circumscribed Sphere Area Ratio; R. sphericity – Ratio sphericity.
